## Supplementary Figures & Tables for "Two mosquito odorant receptors with reciprocal specificity mediated by a single amino acid residue"

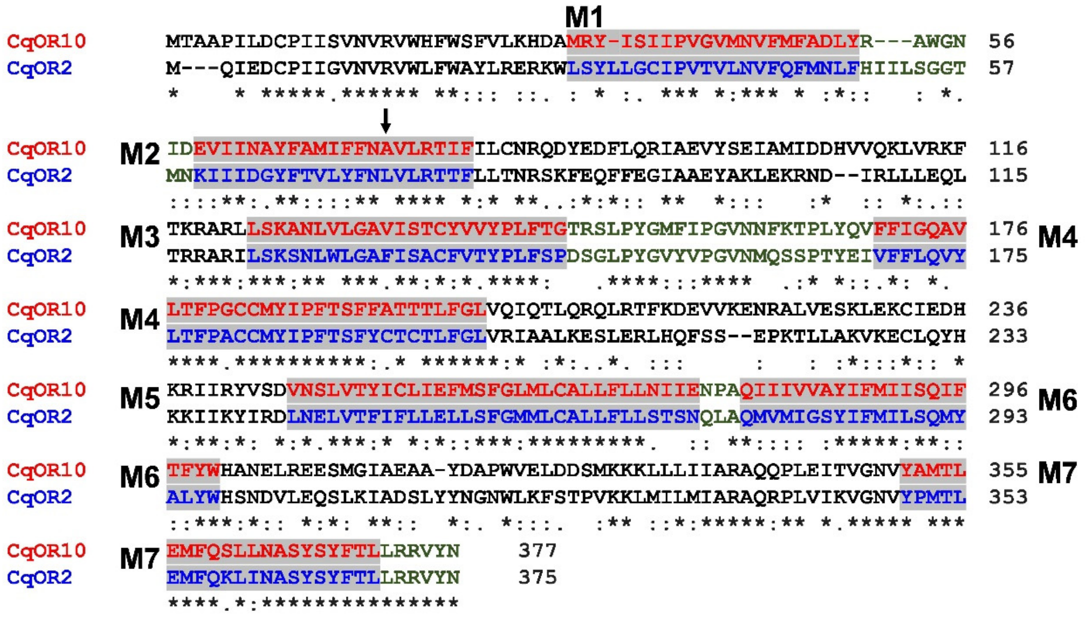


**Figure 1—supplement figure 1.** Alignment of the amino acid sequences of CquiOR10 (abbreviated CqOR10) and CquiOR2 (=CqOR2). OCTOPUS predicted the seven transmembrane domains (M1-M7). The sequences of the N-terminus and intracellular loops are in black, C-terminus and extracellular loops in green, whereas the transmembrane domains of CquiOR10 and CquiOR2 are in red and blue, respectively.

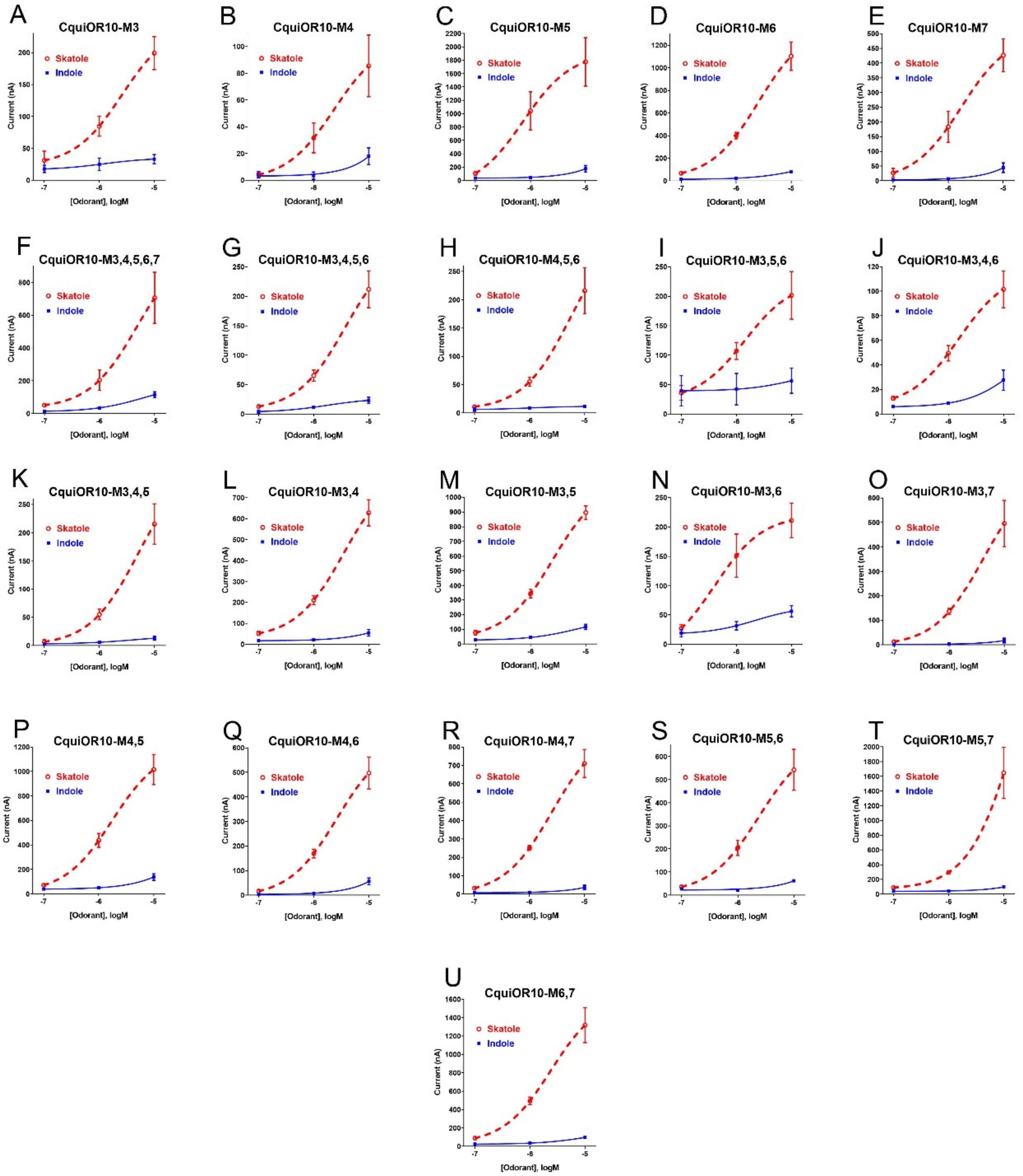


**Figure 1—supplement figure 2.** Concentration-response analysis for activation of wild-type and chimeric ORs by skatole and indole. (A) CquiOR10$^{M3}$; (B) CquiOR10$^{M4}$; (C) CquiOR10$^{M5}$; (D) CquiOR10$^{M6}$; (E) CquiOR10$^{M7}$; (F) CquiOR10$^{M3,4,5,6,7}$; (G) CquiOR10$^{M3,4,5,6}$; (H) CquiOR10$^{M4,5,6}$; (I) CquiOR10$^{M3,5,6}$; (J) CquiOR10$^{M3,4,6}$; (K) CquiOR10$^{M3,4,5}$; (L) CquiOR10$^{M3,4}$; (M) CquiOR10$^{M3,5}$; (N) CquiOR10$^{M3,6,}$; (O) CquiOR10$^{M3,7}$; (P) CquiOR10$^{M4,5}$; (Q) CquiOR10$^{M4,6}$; (R) CquiOR10$^{M4,7}$; (S) CquiOR10$^{M5,6}$; (T) CquiOR10$^{M5,7}$; and (U) CquiOR10$^{M6,7}$. Lines were obtained with nonlinear fit. Bars represent SEM. *n* = 3.

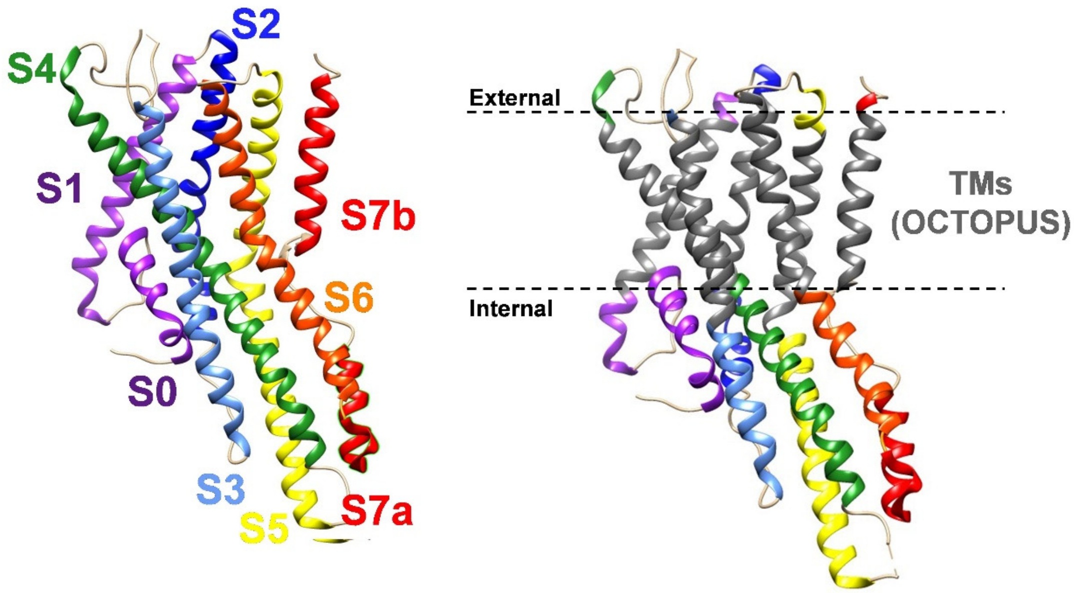


**Figure 1—supplement figure 3.**The Cryo-EM structure of the odorant receptor coreceptor, AbakOrco (PDB, 6C70) generated by UCSF Chimera and displayed in rainbow color code (left). The predicted transmembrane domains (right) are displayed in black. The dashed lines represent the membrane boundaries.

| | 57 … 73 … 116 | Selective to |
|---|---|---|
| CquiOR10 | IDEVIINAYFAMIFFNAVLRTIFILCNRQDYEDFLQRIAEVYSEIAMIDDHVVQKLVRKF | Skatole |
| CquiOR10M$^{2,5,6,7}$ | IDKIIIDGYFTVLYFNLVLRTTFILCNRQDYEDFLQRIAEVYSEIAMIDDHVVQKLVRKF | Indole |
| | Outer / Mid / Inner | |
| CquiOR10M$^{2,5,6,7}$_Outer | IDEVIINAYFTVLYFNLVLRTTFILCNRQDYEDFLQRIAEVYSEIAMIDDHVVQKLVRKF | Indole |
| CquiOR10M$^{2,5,6,7}$_Mid+Inner | IDKIIIDGYFAMIFFNAVLRTIFILCNRQDYEDFLQRIAEVYSEIAMIDDHVVQKLVRKF | Skatole |
| CquiOR10M$^{2,5,6,7}$_Inner | IDKIIIDGYFTVLYFNAVLRTIFILCNRQDYEDFLQRIAEVYSEIAMIDDHVVQKLVRKF | Skatole |
| CquiOR10M$^{2,5,6,7}$_T78I | IDKIIIDGYFTVLYFNLVLRTIFILCNRQDYEDFLQRIAEVYSEIAMIDDHVVQKLVRKF | Indole |
| CquiOR10M$^{2,5,6,7}$_L73A | IDKIIIDGYFTVLYFNAVLRTTFILCNRQDYEDFLQRIAEVYSEIAMIDDHVVQKLVRKF | Skatole |

**Figure 2—supplement figure 1.** Partial sequences of CquiOR10 and chimeric ORs highlighting transmembrane domain-2 (TM2). The two last residues of the extracellular loop-1 (Ile-57 and Asp-58) appear in the N-terminus. The TM2 was divided into the arbitrary segments outer, middle (mid), and inner to identify specificity determinants.

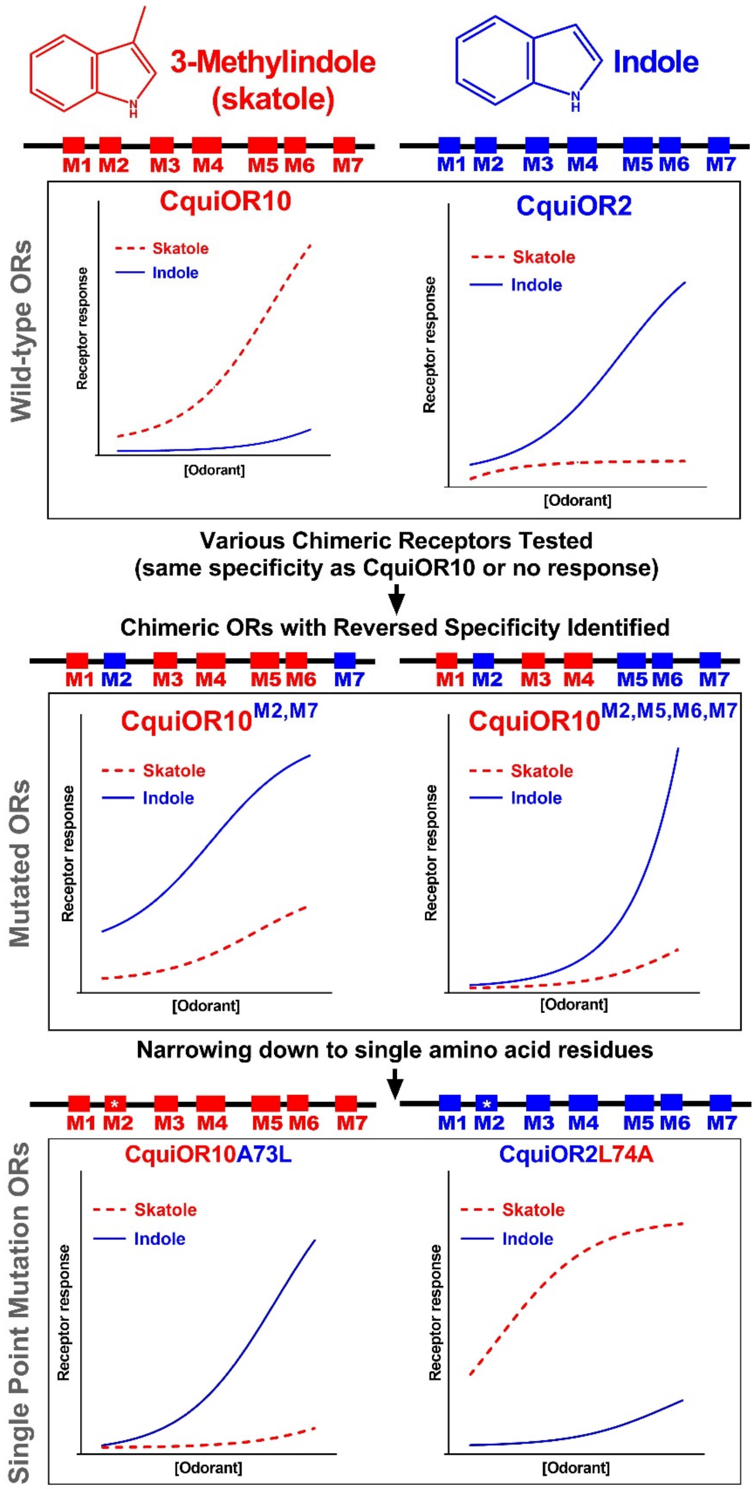


**Figure 2—supplement figure 2.** Graphical representation summarizing the reciprocal specificity of CquiOR10 and CquiOR2 mediated by a single amino acid residue.

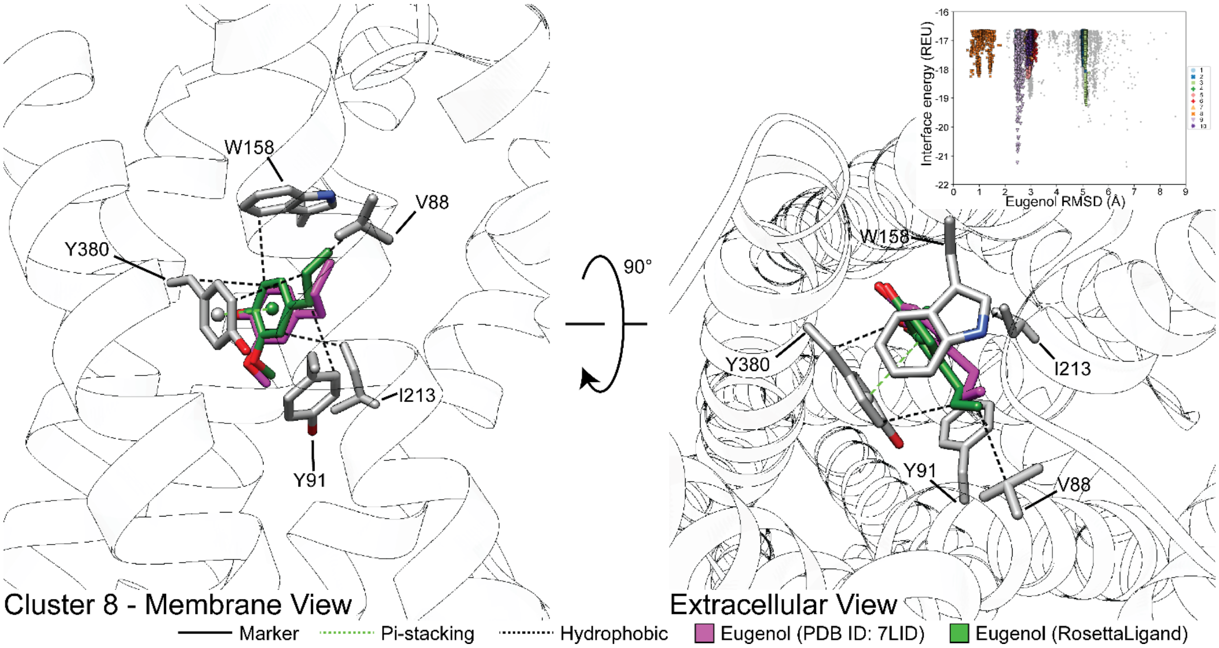


**Figure 5—supplement figure 1.** Representative model of docked eugenol with MhraOR5 using RosettaLigand matches experimental structure. The model represented (forest green) is the lowest interface-energy pose from cluster 8. From cluster 8, the lowest RMSD pose was 0.55 Å relative to the experimental coordinates of eugenol from PDB ID: 7LID (magenta). Atoms that are not eugenol carbon atoms are color coded by atom type: carbon (grey), nitrogen (dark blue), and oxygen (red). The inset represents the clustering of the 10,000 lowest interface-energy models, plotted by Rosetta interface-energy (Rosetta Energy Units) and by ligand RMSD (Å). The legend within the inset represents the top 10 most frequent clusters, sorted by largest to smallest (various color/shape combinations) The grey circles represent structure models that were not clustered by the algorithm.

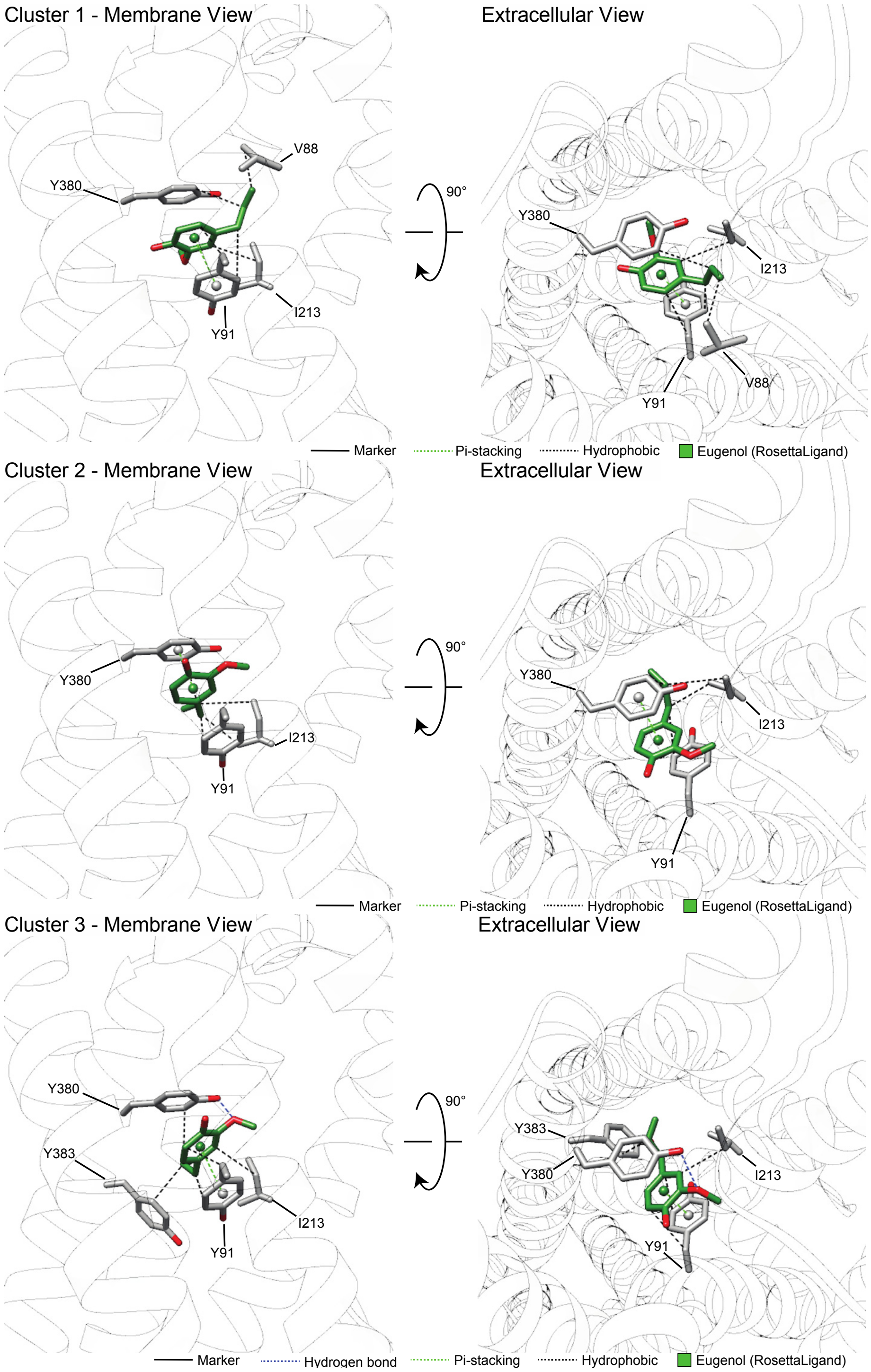

Cluster 1 - Membrane View
Extracellular View
V88
Y380
I213
Y91
90°
Y380
I213
Y91
V88
Marker
Pi-stacking
Hydrophobic
Eugenol (RosettaLigand)
Cluster 2 - Membrane View
Extracellular View
Y380
I213
Y91
90°
Y380
I213
Y91
Marker
Pi-stacking
Hydrophobic
Eugenol (RosettaLigand)
Cluster 3 - Membrane View
Extracellular View
Y380
Y383
I213
Y91
90°
Y383
Y380
I213
Y91
Marker
Hydrogen bond
Pi-stacking
Hydrophobic
Eugenol (RosettaLigand)

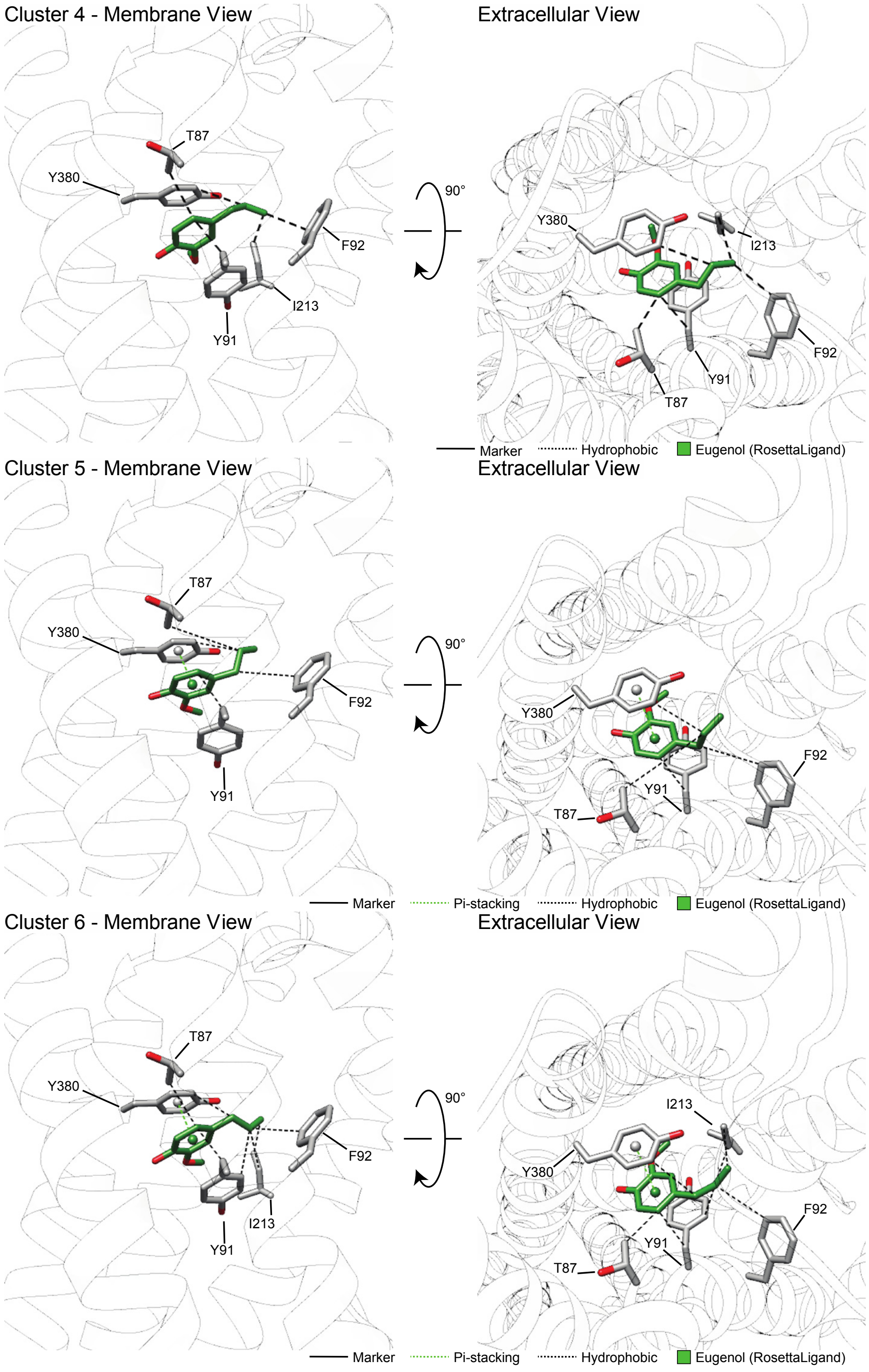
Cluster 4 - Membrane View
Extracellular View
T87
Y380
F92
I213
Y91
90°
Marker
Hydrophobic
Eugenol (RosettaLigand)
Cluster 5 - Membrane View
Extracellular View
Pi-stacking
Cluster 6 - Membrane View

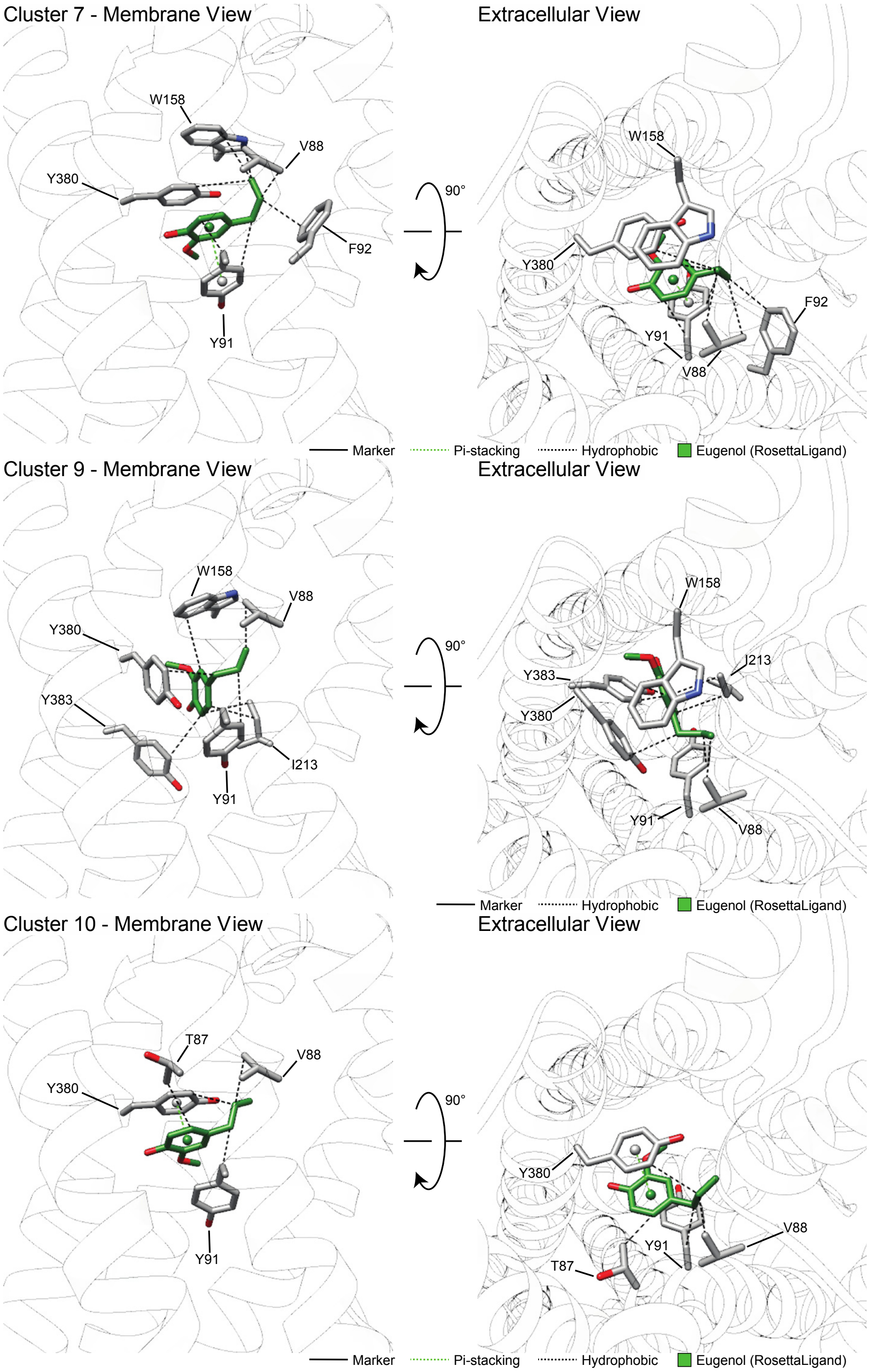


**Figure 5—supplement figure 2.** Clusters 1-7, 9,10 from RosettaLigand docking of eugenol to MhraOR5 with PLIP analysis. Hydrogen bond and pi-stacking interactions were filtered by previously reported bond distances (Bissantz et al., 2010).

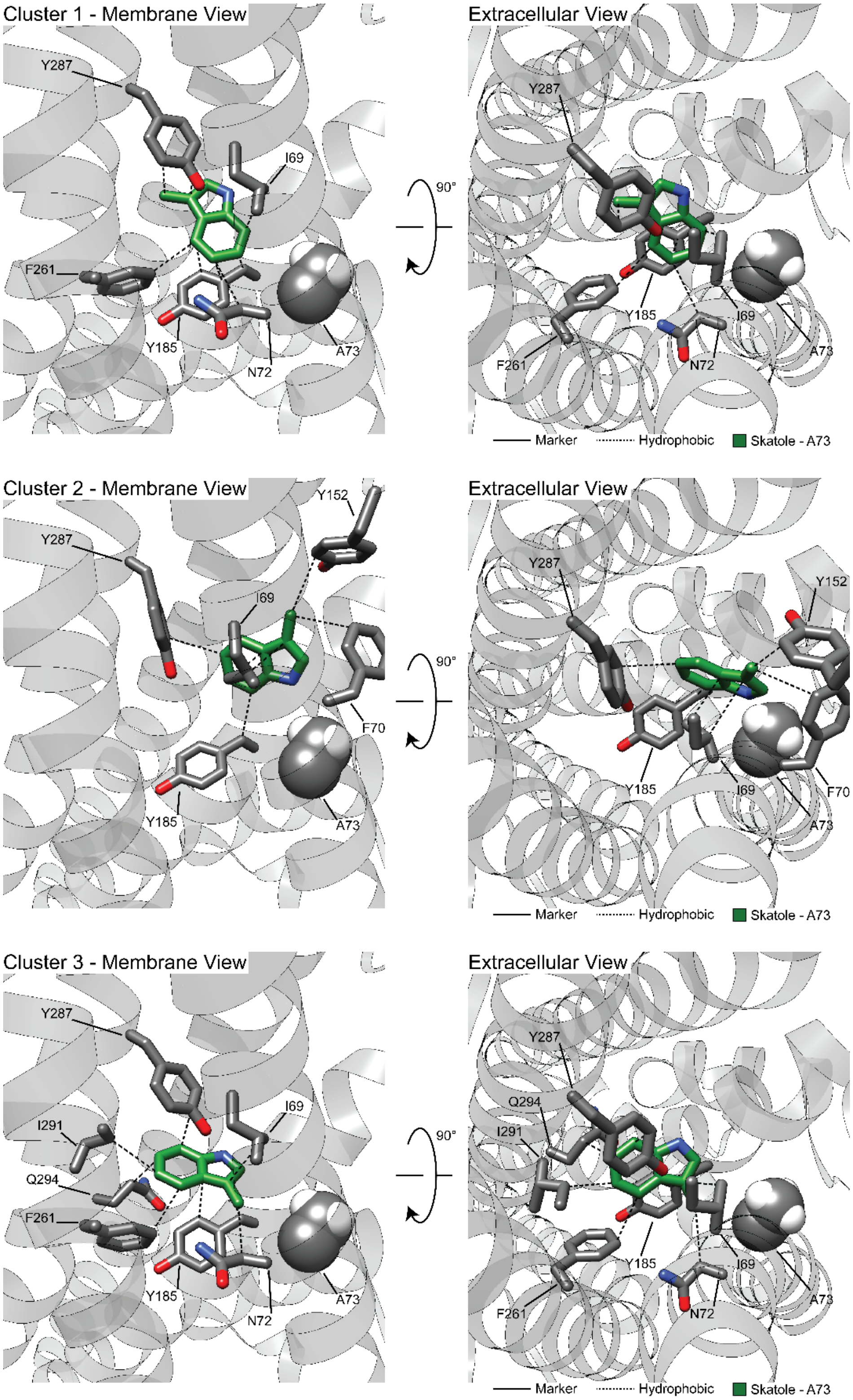
Cluster 1 - Membrane View
Extracellular View
Y287
I69
F261
Y185
N72
A73
90°
Marker
Hydrophobic
Skatole - A73
Cluster 2 - Membrane View
Extracellular View
Y152
Y287
I69
F70
Y185
A73
90°
Marker
Hydrophobic
Skatole - A73
Cluster 3 - Membrane View
Extracellular View
Y287
I69
I291
Q294
F261
Y185
N72
A73
90°
Marker
Hydrophobic
Skatole - A73

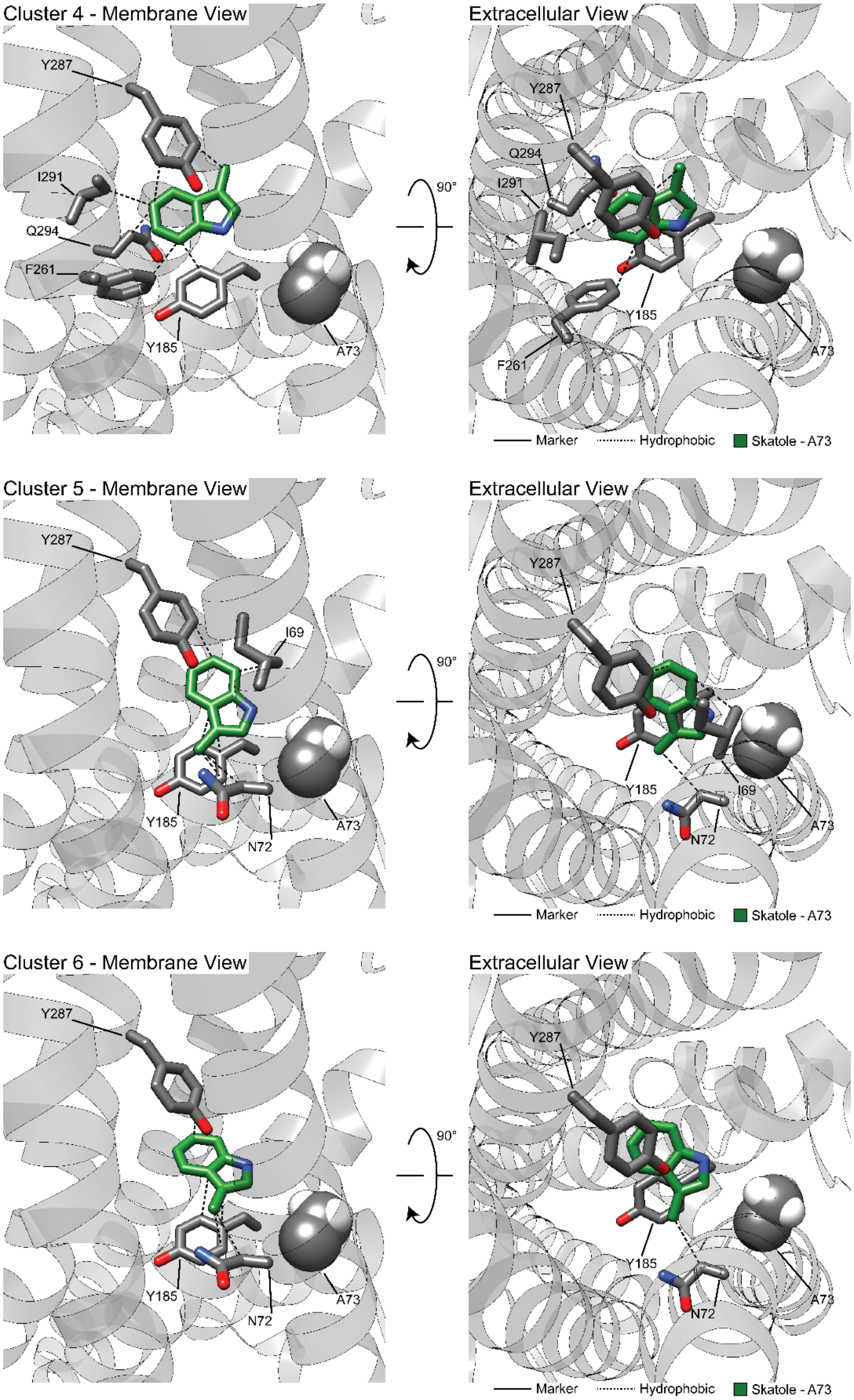

Cluster 4 - Membrane View
Extracellular View
Y287
I291
Q294
F261
Y185
A73
90°
Marker
Hydrophobic
Skatole - A73
Cluster 5 - Membrane View
Extracellular View
I69
N72
Cluster 6 - Membrane View
Extracellular View

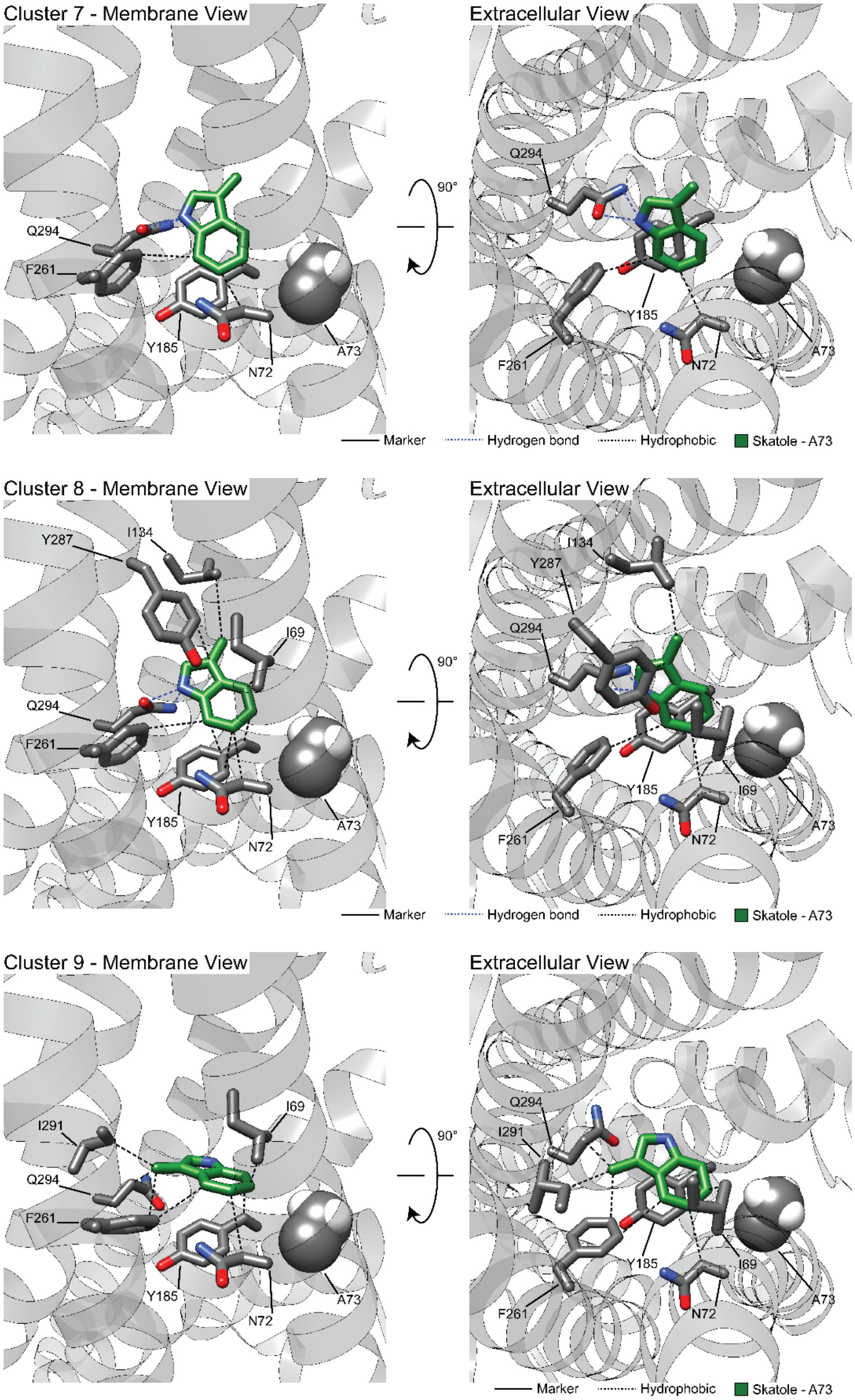

Cluster 7 - Membrane View
Extracellular View
Q294
F261
Y185
N72
A73
90°
Marker
Hydrogen bond
Hydrophobic
Skatole - A73
Cluster 8 - Membrane View
Extracellular View
Y287
I134
I69
Q294
F261
Y185
N72
A73
90°
Marker
Hydrogen bond
Hydrophobic
Skatole - A73
Cluster 9 - Membrane View
Extracellular View
I291
Q294
F261
I69
Y185
N72
A73
90°
Marker
Hydrophobic
Skatole - A73

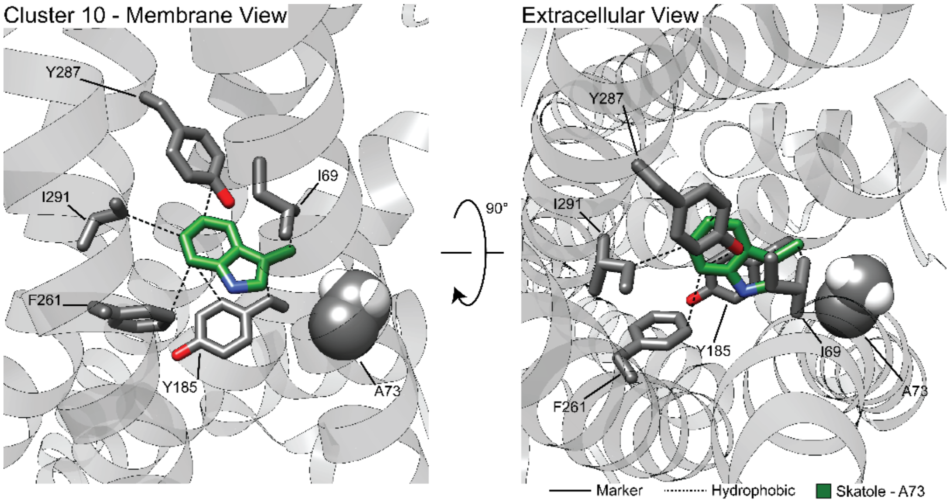


**Figure 6—supplement figure 1A.** Clusters 1-10 from RosettaLigand docking of skatole to CquiOR10 with PLIP analysis. Hydrogen bond and pi-stacking interactions were filtered by previously reported bond distances (Bissantz et al., 2010).

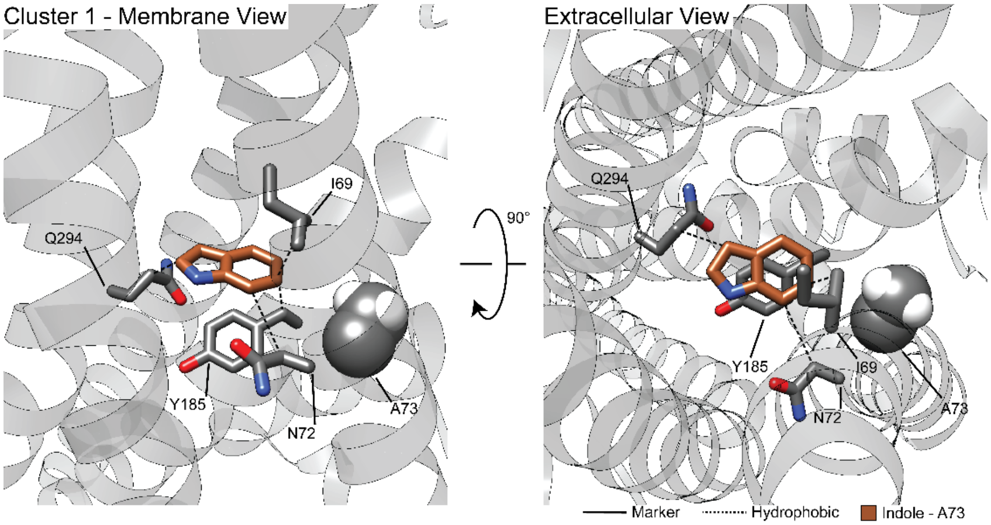

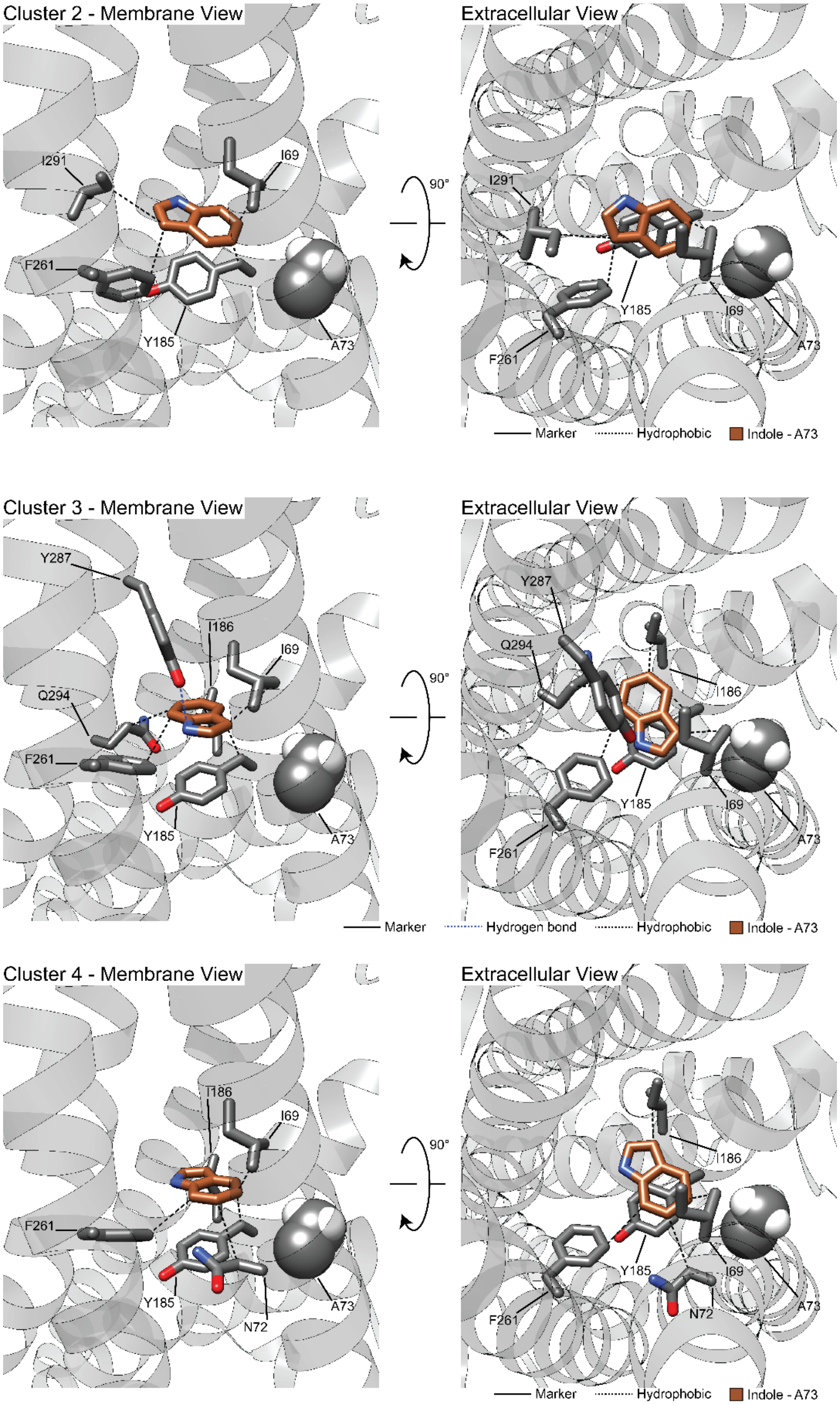
Cluster 2 - Membrane View
I291
I69
F261
Y185
A73
90°
Extracellular View
I291
Y185
I69
F261
A73
Marker
Hydrophobic
Indole - A73
Cluster 3 - Membrane View
Y287
I186
I69
Q294
F261
Y185
A73
90°
Extracellular View
Y287
Q294
I186
Y185
I69
F261
A73
Marker
Hydrogen bond
Hydrophobic
Indole - A73
Cluster 4 - Membrane View
I186
I69
F261
Y185
N72
A73
90°
Extracellular View
I186
Y185
I69
N72
F261
A73
Marker
Hydrophobic
Indole - A73

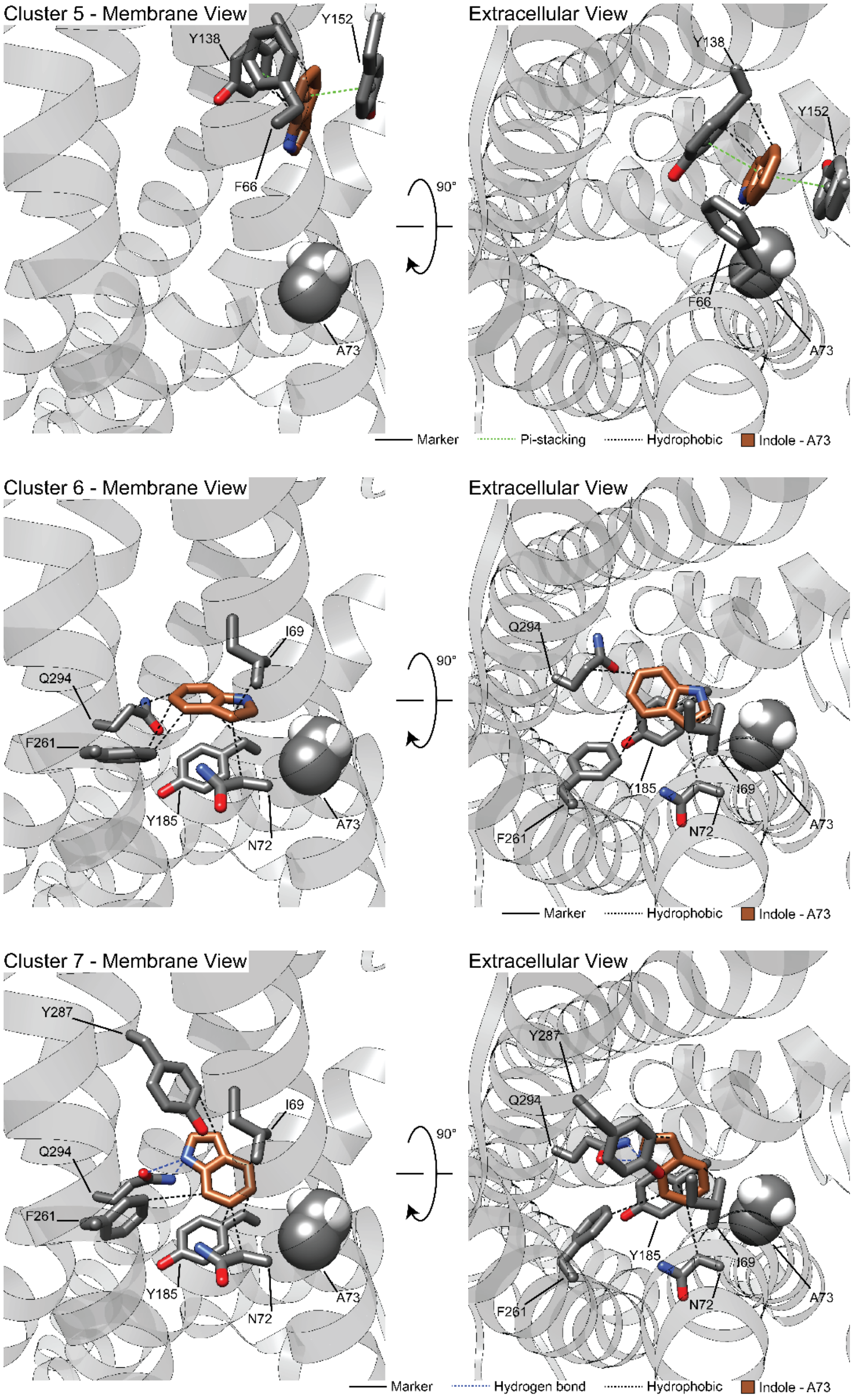
Cluster 5 - Membrane View
Y138
Y152
F66
A73
Extracellular View
90°
Y138
Y152
F66
A73
Marker
Pi-stacking
Hydrophobic
Indole - A73
Cluster 6 - Membrane View
I69
Q294
F261
Y185
N72
A73
Extracellular View
90°
Q294
Y185
I69
F261
N72
A73
Marker
Hydrophobic
Indole - A73
Cluster 7 - Membrane View
Y287
I69
Q294
F261
Y185
N72
A73
Extracellular View
90°
Y287
Q294
Y185
I69
F261
N72
A73
Marker
Hydrogen bond
Hydrophobic
Indole - A73

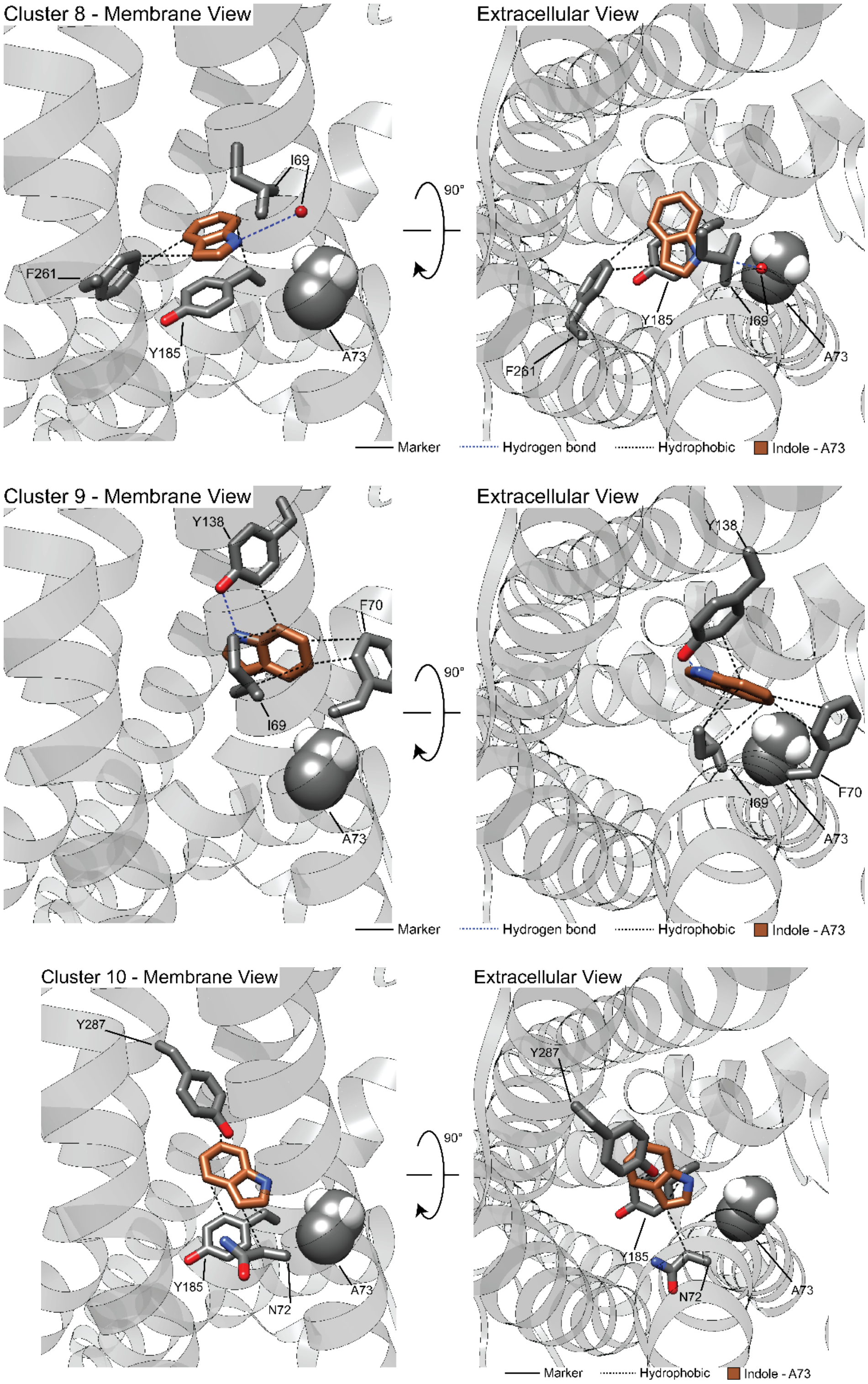


**Figure 6—supplement figure 1B.** Clusters 1-10 from RosettaLigand docking of indole to CquiOR10 with PLIP analysis. Hydrogen bond and pi-stacking interactions were filtered by previously reported bond distances (Bissantz et al., 2010).

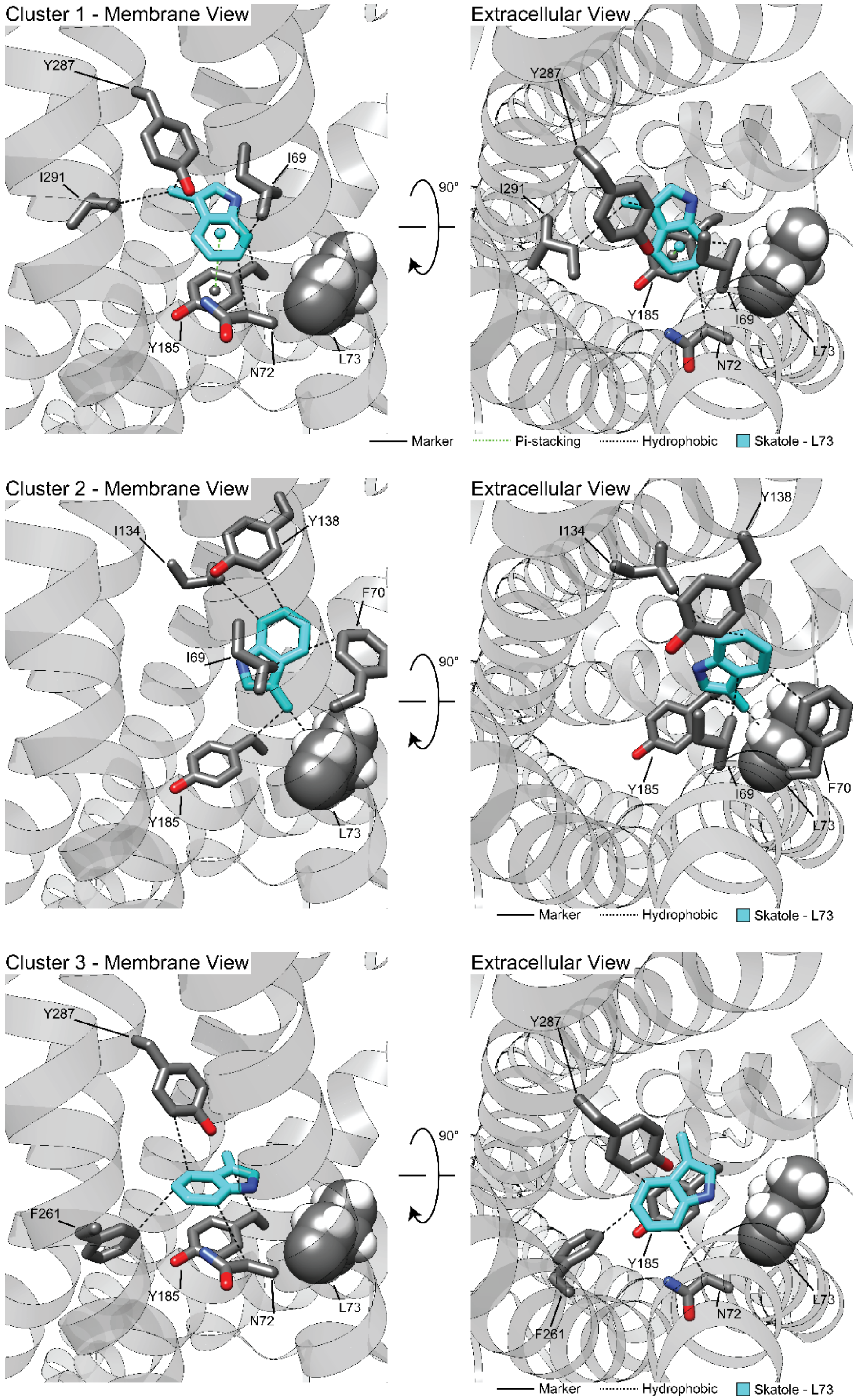
Cluster 1 - Membrane View
Extracellular View
Y287
I291
I69
Y185
N72
L73
90°
Marker
Pi-stacking
Hydrophobic
Skatole - L73
Cluster 2 - Membrane View
Extracellular View
I134
Y138
F70
I69
Y185
L73
90°
Marker
Hydrophobic
Skatole - L73
Cluster 3 - Membrane View
Extracellular View
Y287
F261
Y185
N72
L73
90°
Marker
Hydrophobic
Skatole - L73

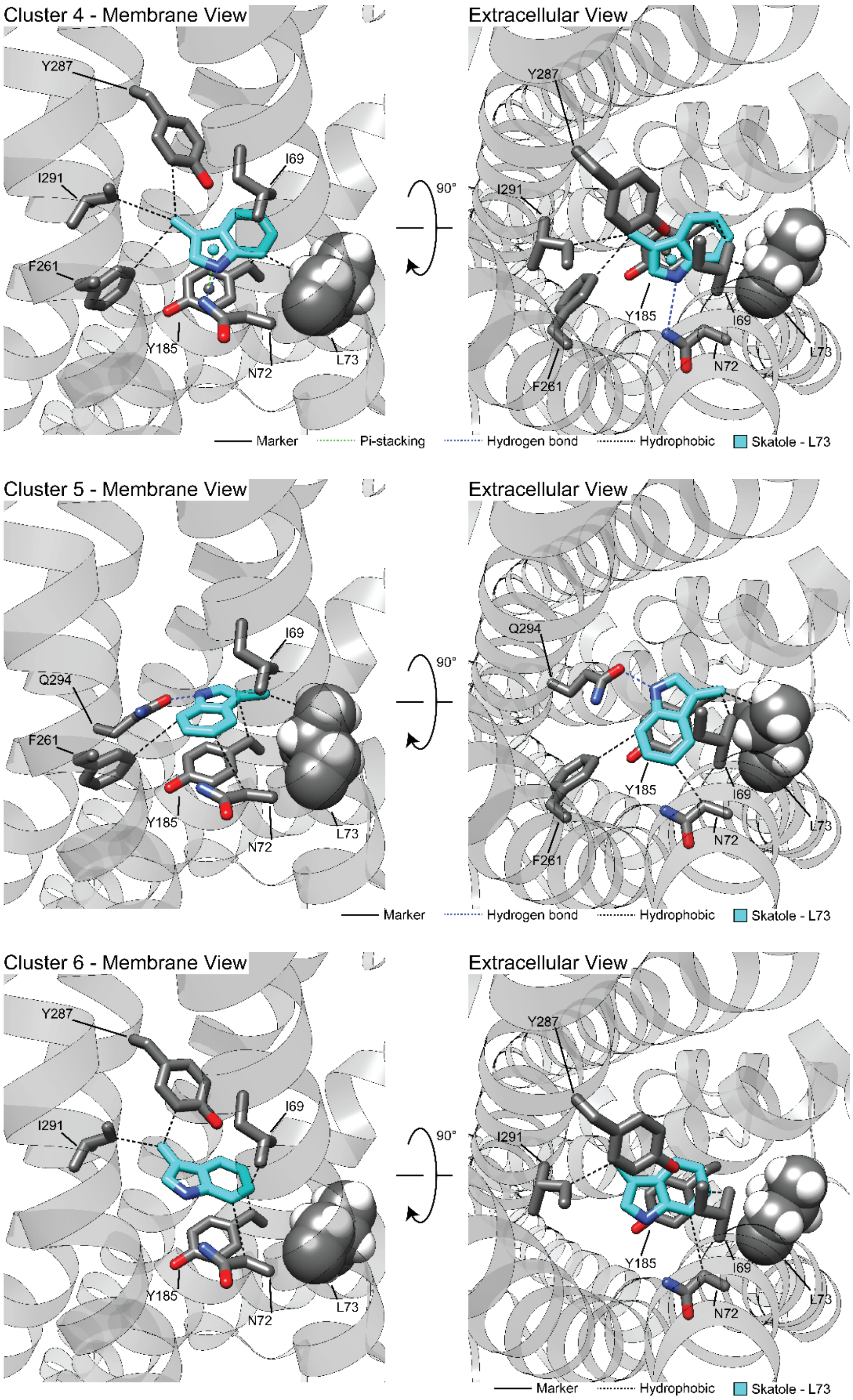
Cluster 4 - Membrane View
Extracellular View
Y287
I291
I69
F261
Y185
N72
L73
90°
Marker
Pi-stacking
Hydrogen bond
Hydrophobic
Skatole - L73
Cluster 5 - Membrane View
Extracellular View
Q294
I69
F261
Y185
N72
L73
90°
Marker
Hydrogen bond
Hydrophobic
Skatole - L73
Cluster 6 - Membrane View
Extracellular View
Y287
I291
I69
Y185
N72
L73
90°
Marker
Hydrophobic
Skatole - L73

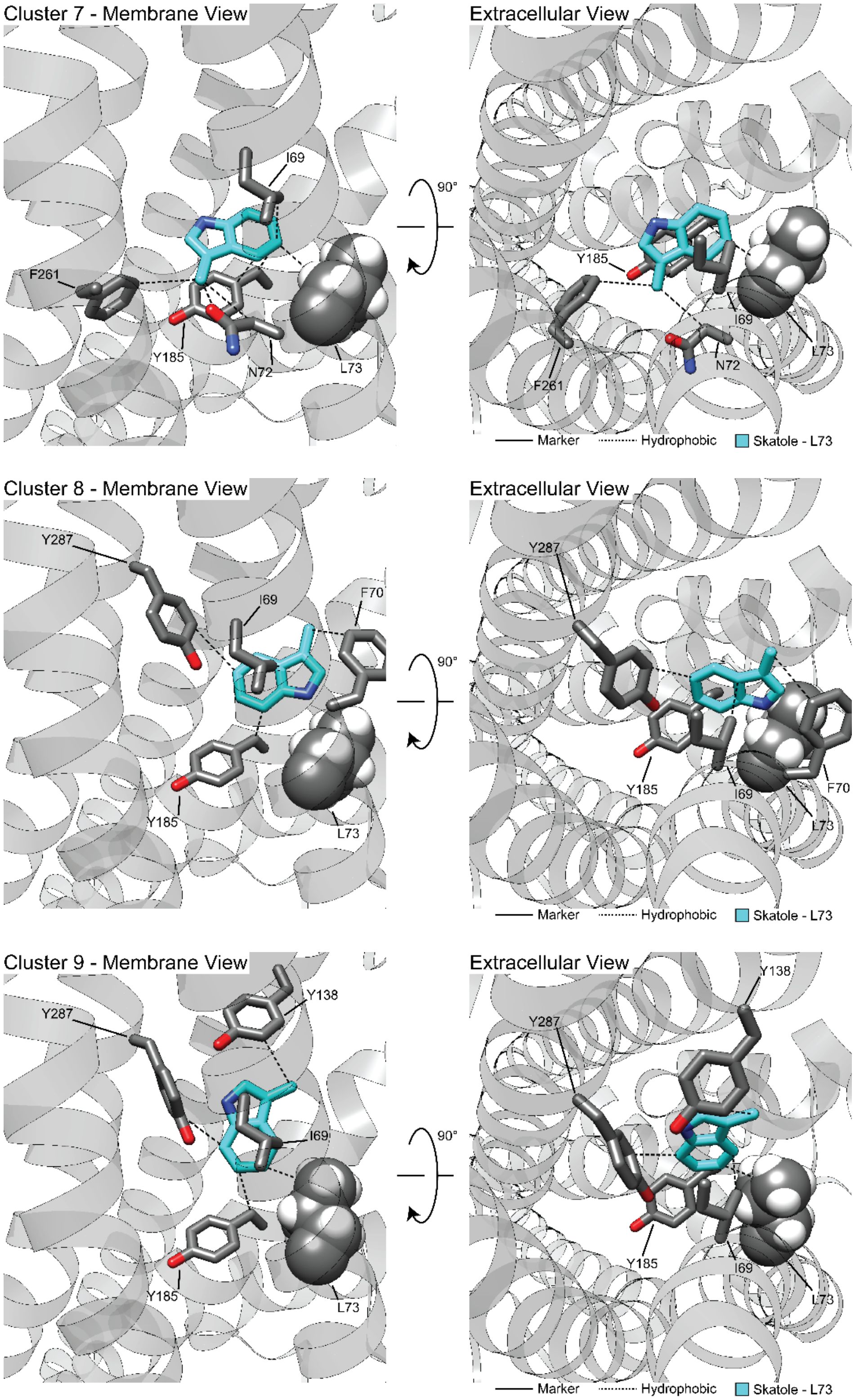
Cluster 7 - Membrane View
Extracellular View
I69
F261
Y185
N72
L73
90°
Marker
Hydrophobic
Skatole - L73
Cluster 8 - Membrane View
Y287
F70
Cluster 9 - Membrane View
Y138

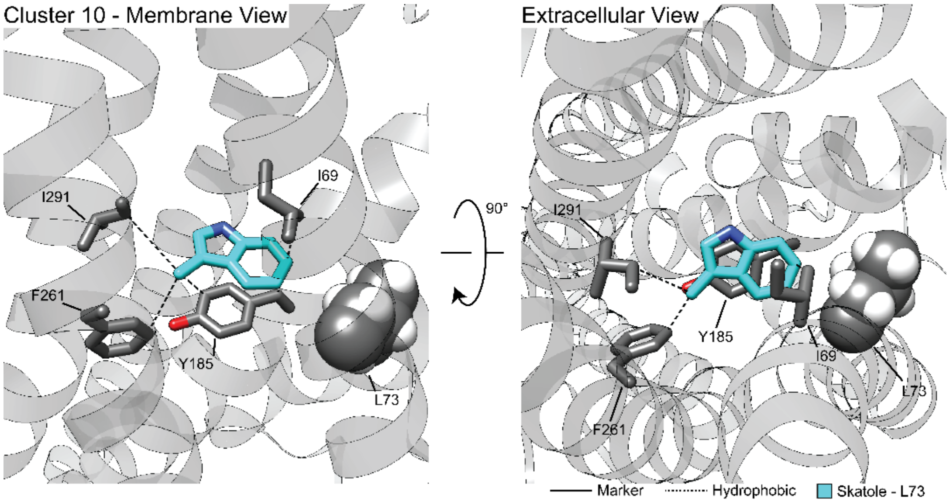


**Figure 6—supplement figure 1C.** Clusters 1-10 from RosettaLigand docking of skatole to CquiOR10A73L with PLIP analysis. Hydrogen bond and pi-stacking interactions were filtered by previously reported bond distances (Bissantz et al., 2010).

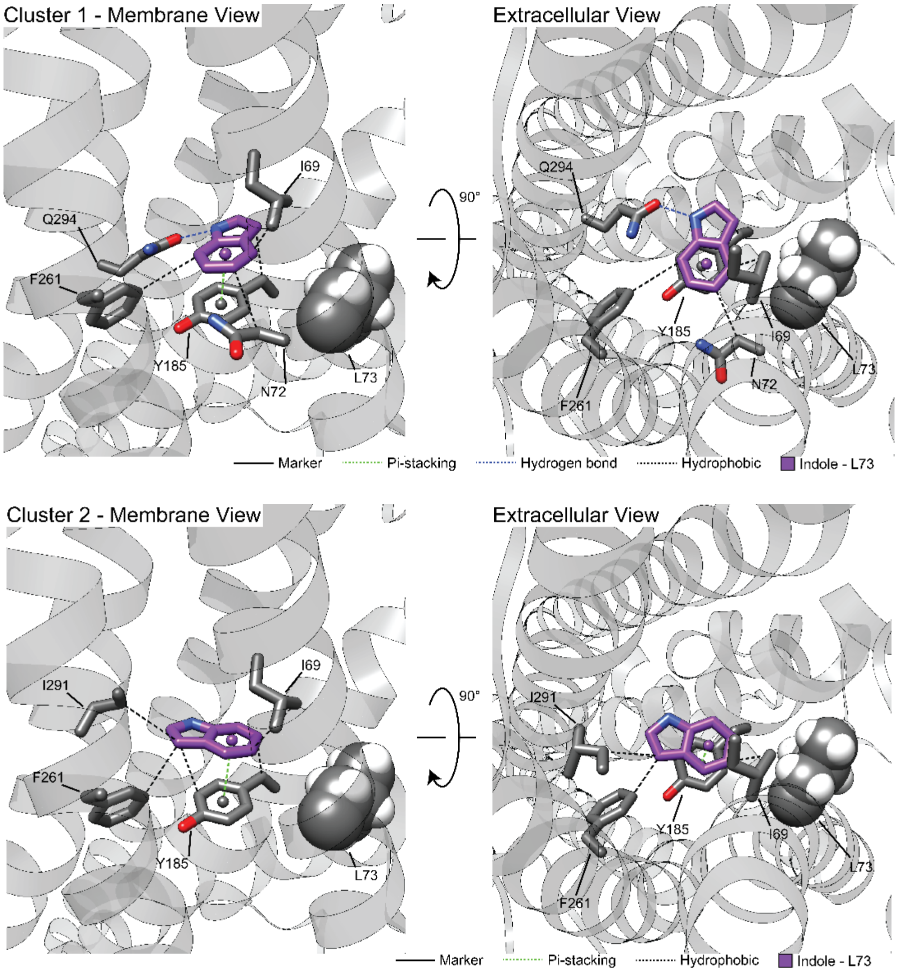

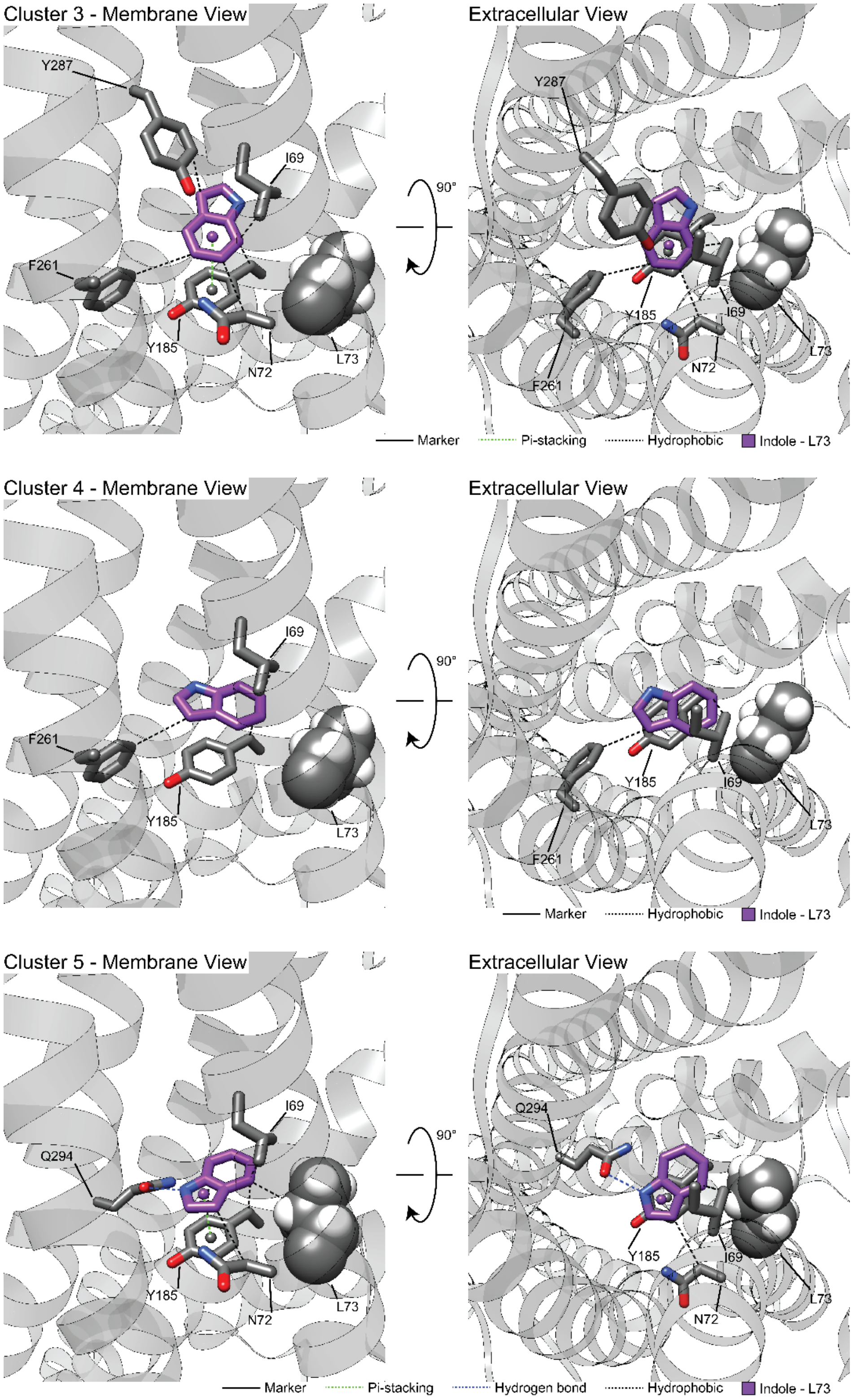
Cluster 3 - Membrane View
Extracellular View
Y287
I69
F261
Y185
N72
L73
90°
Marker
Pi-stacking
Hydrophobic
Indole - L73
Cluster 4 - Membrane View
Extracellular View
Hydrogen bond
Cluster 5 - Membrane View
Q294

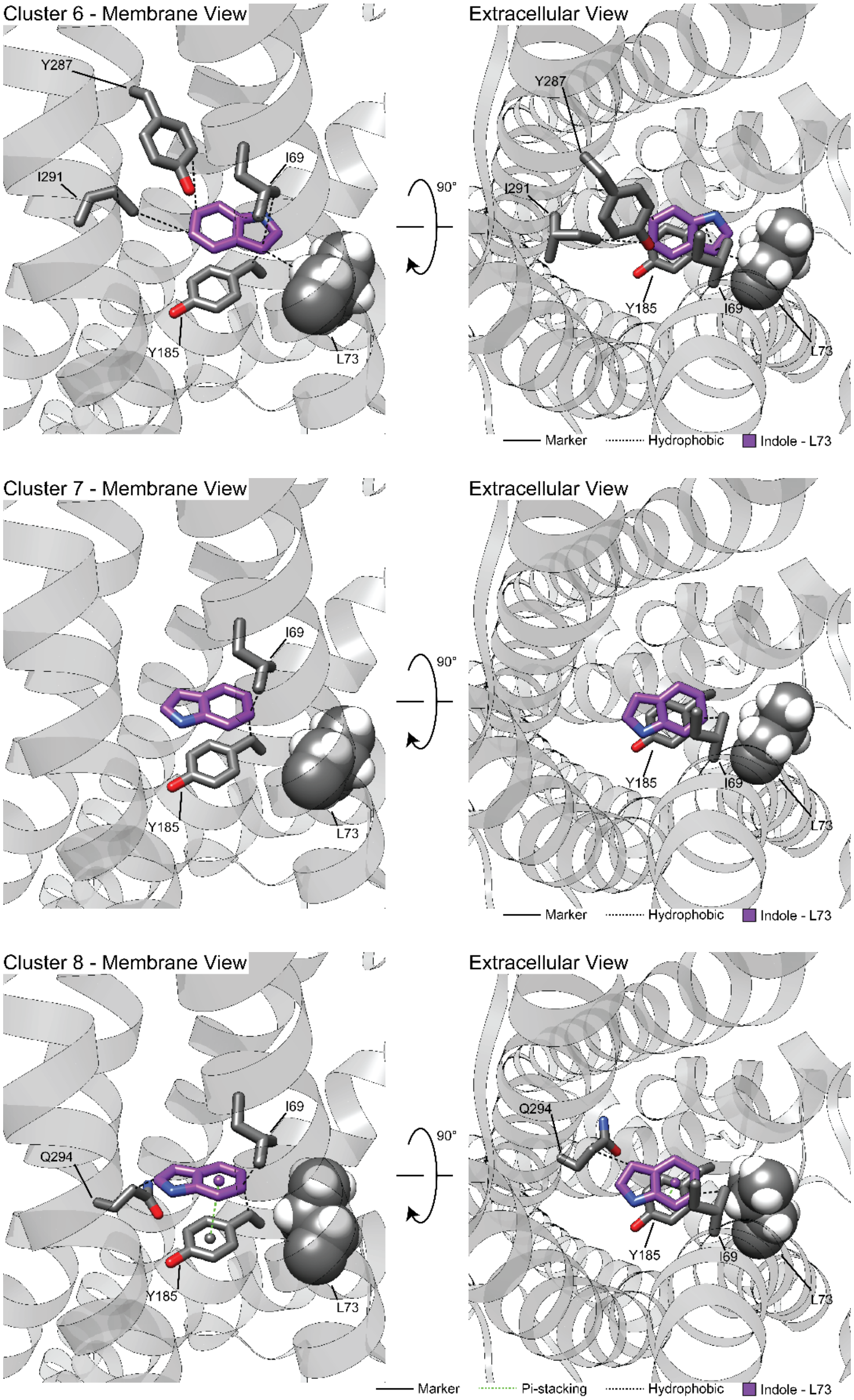

Cluster 6 - Membrane View
Extracellular View
Y287
I69
I291
Y185
L73
90°
Marker
Hydrophobic
Indole - L73
Cluster 7 - Membrane View
Extracellular View
I69
Y185
L73
90°
Marker
Hydrophobic
Indole - L73
Cluster 8 - Membrane View
Extracellular View
I69
Q294
Y185
L73
90°
Marker
Pi-stacking
Hydrophobic
Indole - L73

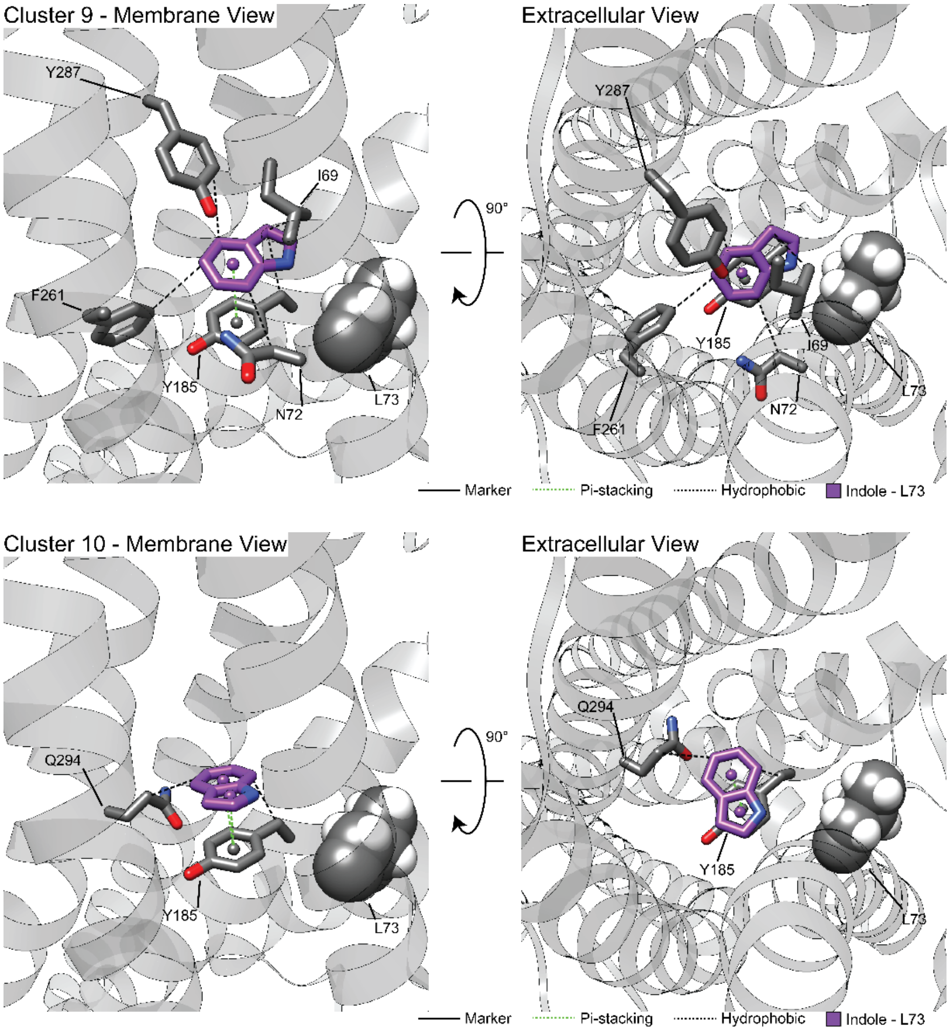


**Figure 6—supplement figure 1D.** Clusters 1-10 from RosettaLigand docking of indole to CquiOR10A73L with PLIP analysis. Hydrogen bond and pi-stacking interactions were filtered by previously reported bond distances (Bissantz et al., 2010).

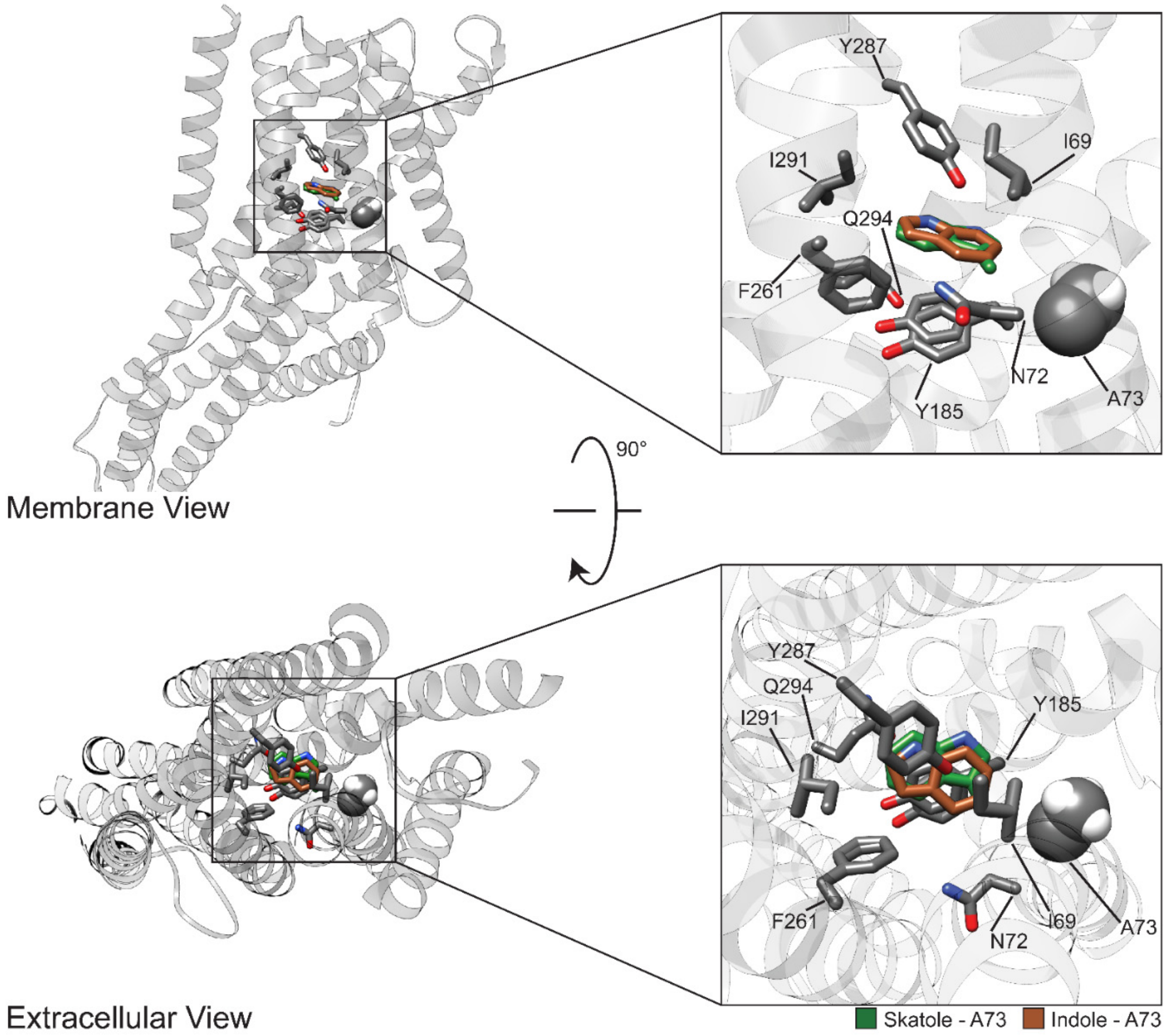


**Figure 6—supplement figure 2A.** CquiOR10 in complex with skatole (forest green) and indole (brown). The lowest interface energy pose's position from the top ten clusters is shown. The entire homomer used in docking (left) and the zoomed-in ligand position with interacting residues (right). A73 shown as spheres.

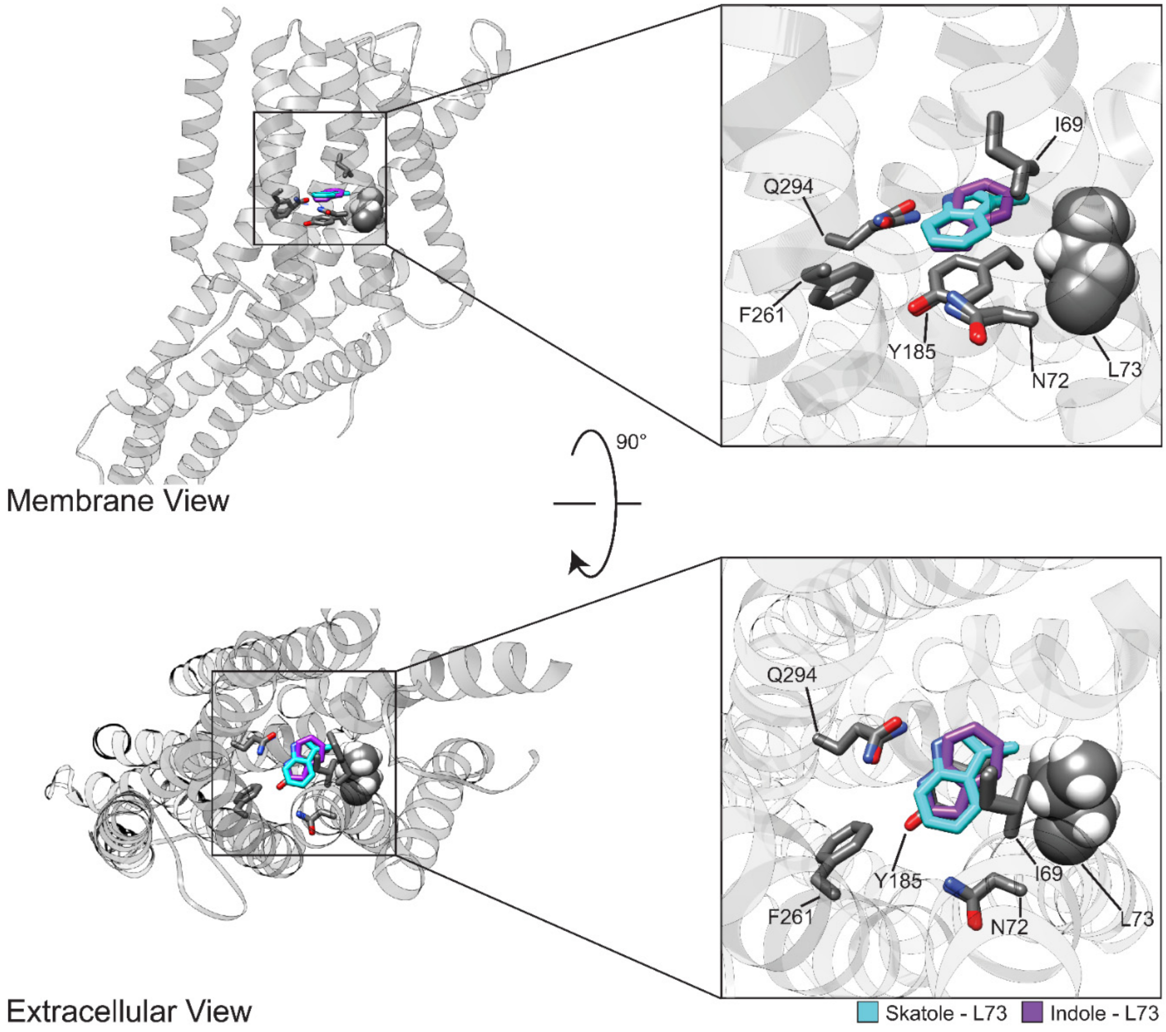


**Figure 6—supplement figure 2B.** CquiOR10-A73L in complex with skatole (light blue) and indole (purple). The lowest interface energy pose's position from the top ten clusters is shown. The entire homomer used in docking (left) and the zoomed-in ligand position with interacting residues (right). L73 shown as spheres.

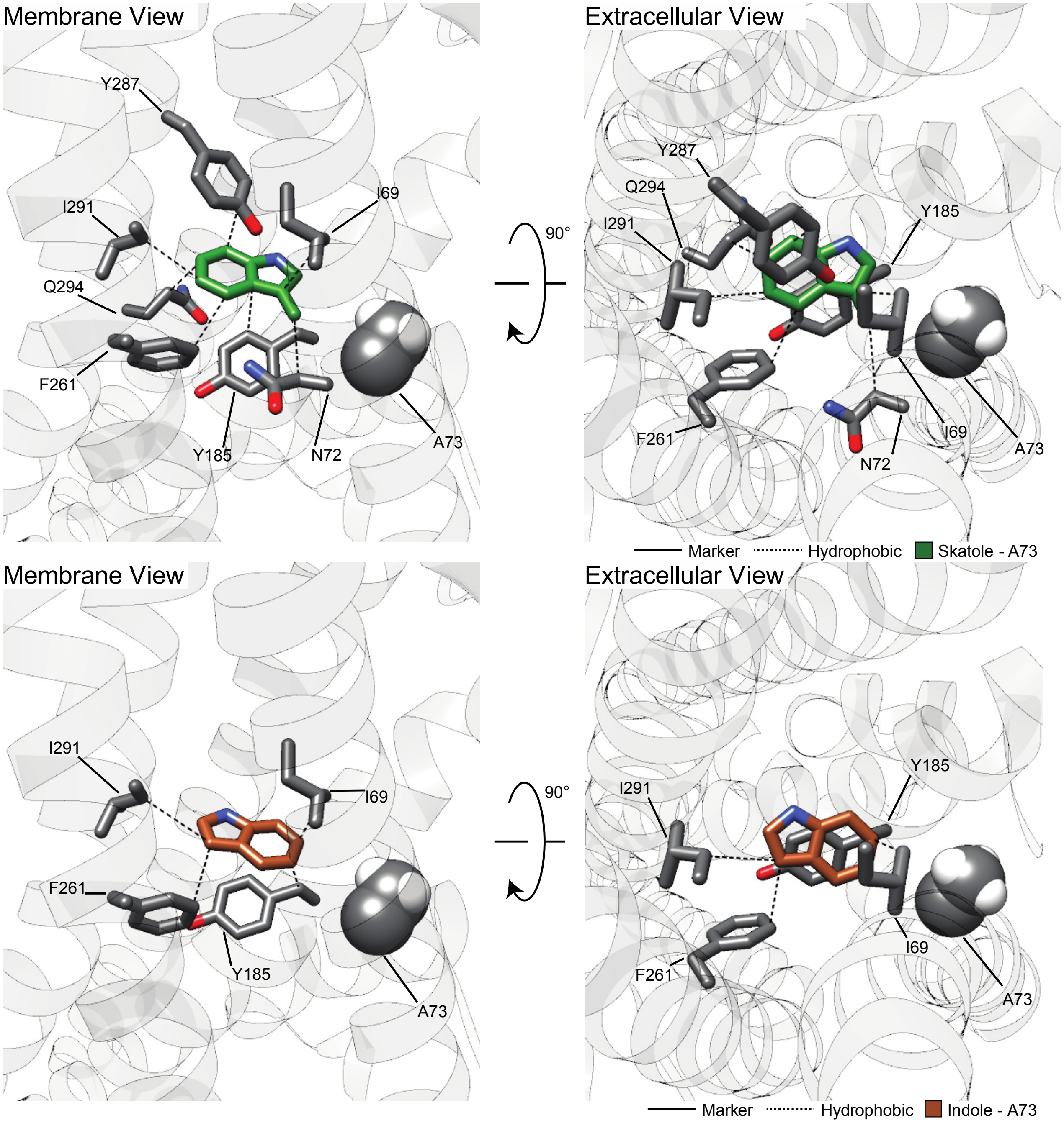


**Figure 6—supplement figure 3A.** Representative RosettaLigand docking of skatole and indole to CquiOR10 with PLIP analysis. The representative is the lowest interface-scoring energy model from the 10 most frequent clusters of each test case. Hydrogen bond and pi-stacking interactions were filtered by previously reported bond distances (Bissantz et al., 2010).

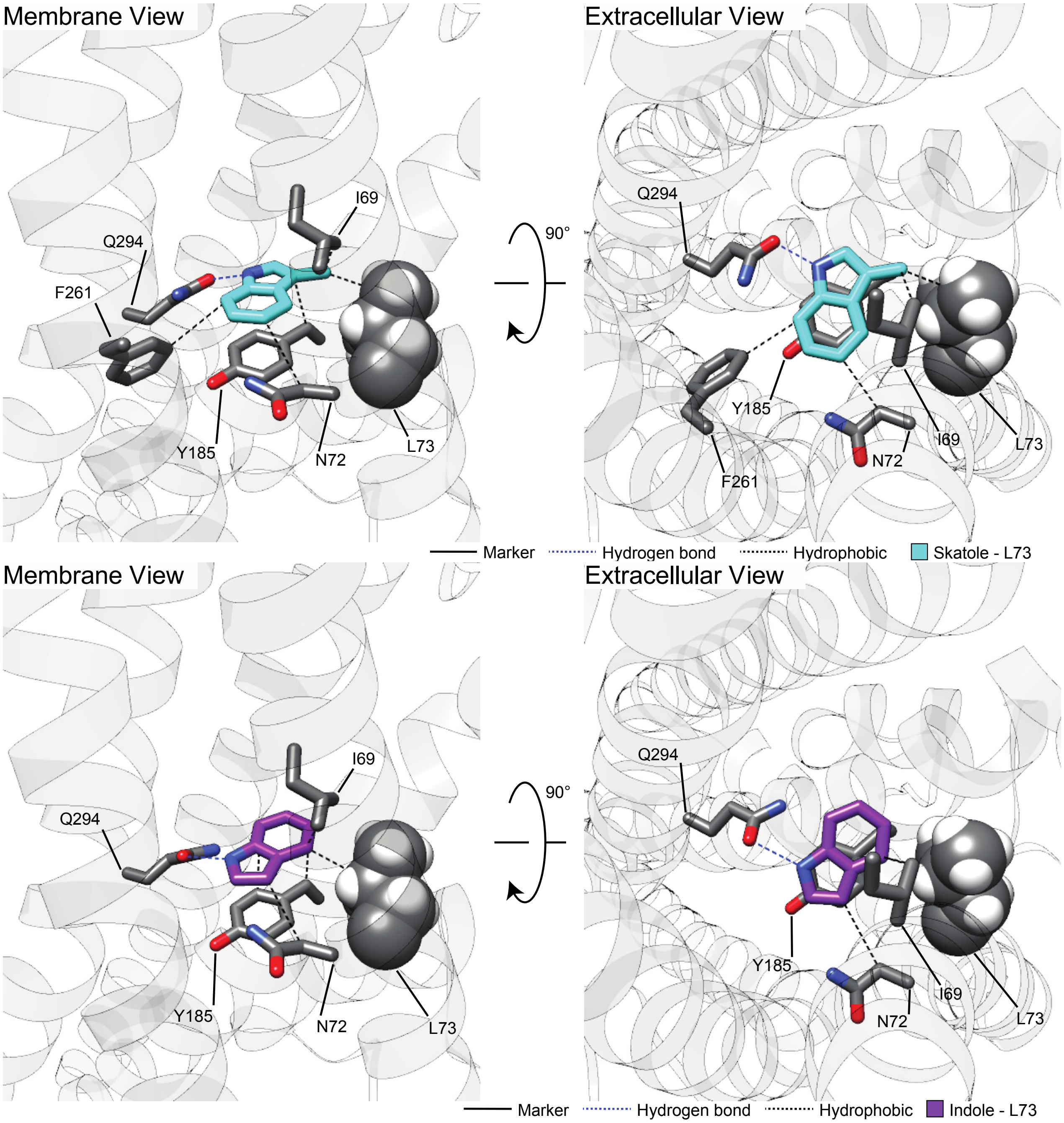


**Figure 6—supplement figure 3B.** Representative RosettaLigand docking of skatole and indole to CquiOR10A73L with PLIP analysis. The representative is the lowest interface-scoring energy model from the 10 most frequent clusters of each test case. Hydrogen bond and pi-stacking interactions were filtered by previously reported bond distances (Bissantz et al., 2010).

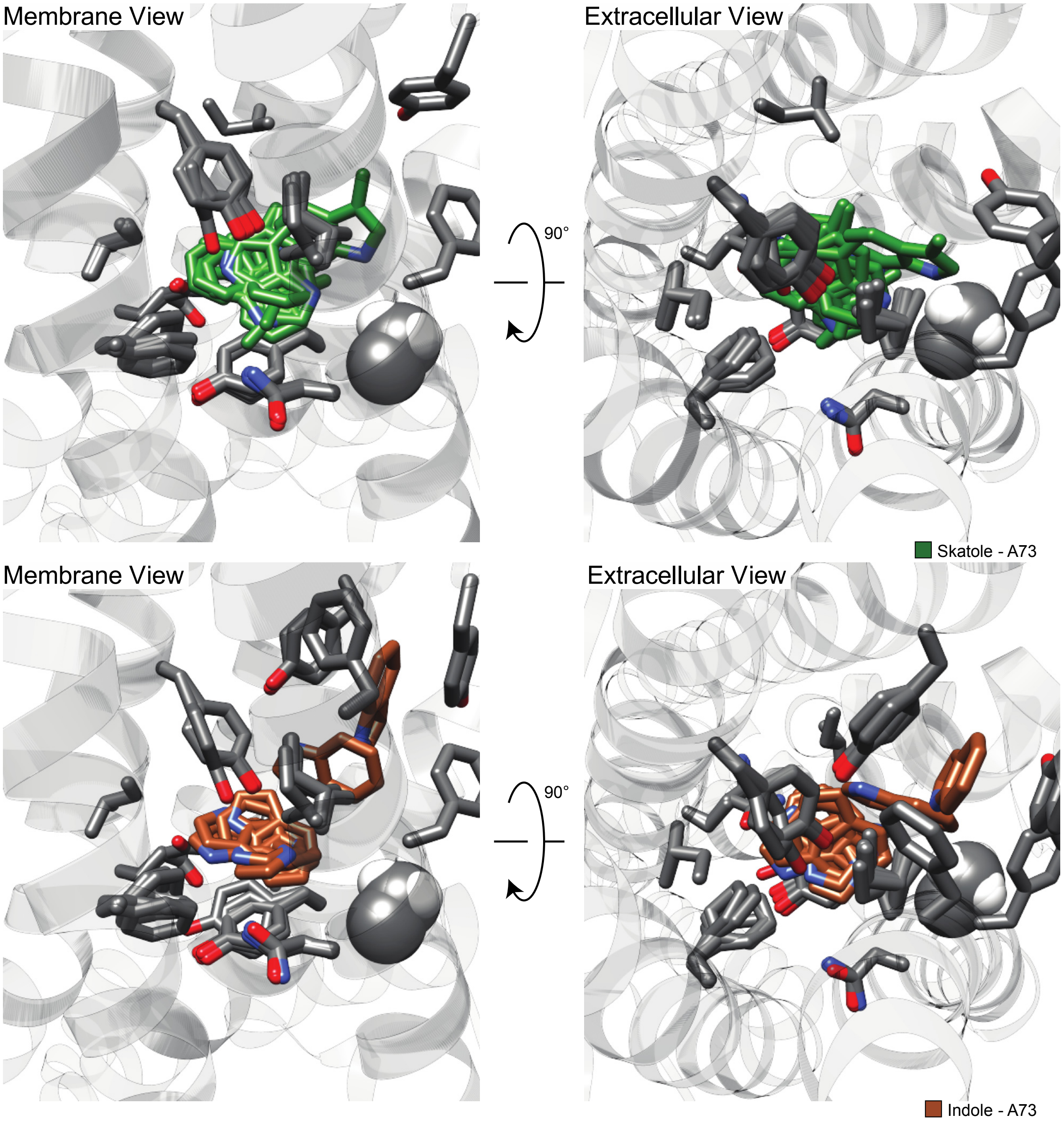


**Figure 6—supplement figure 4A.** Superposition of all clusters of skatole and indole docked to CquiOR10.

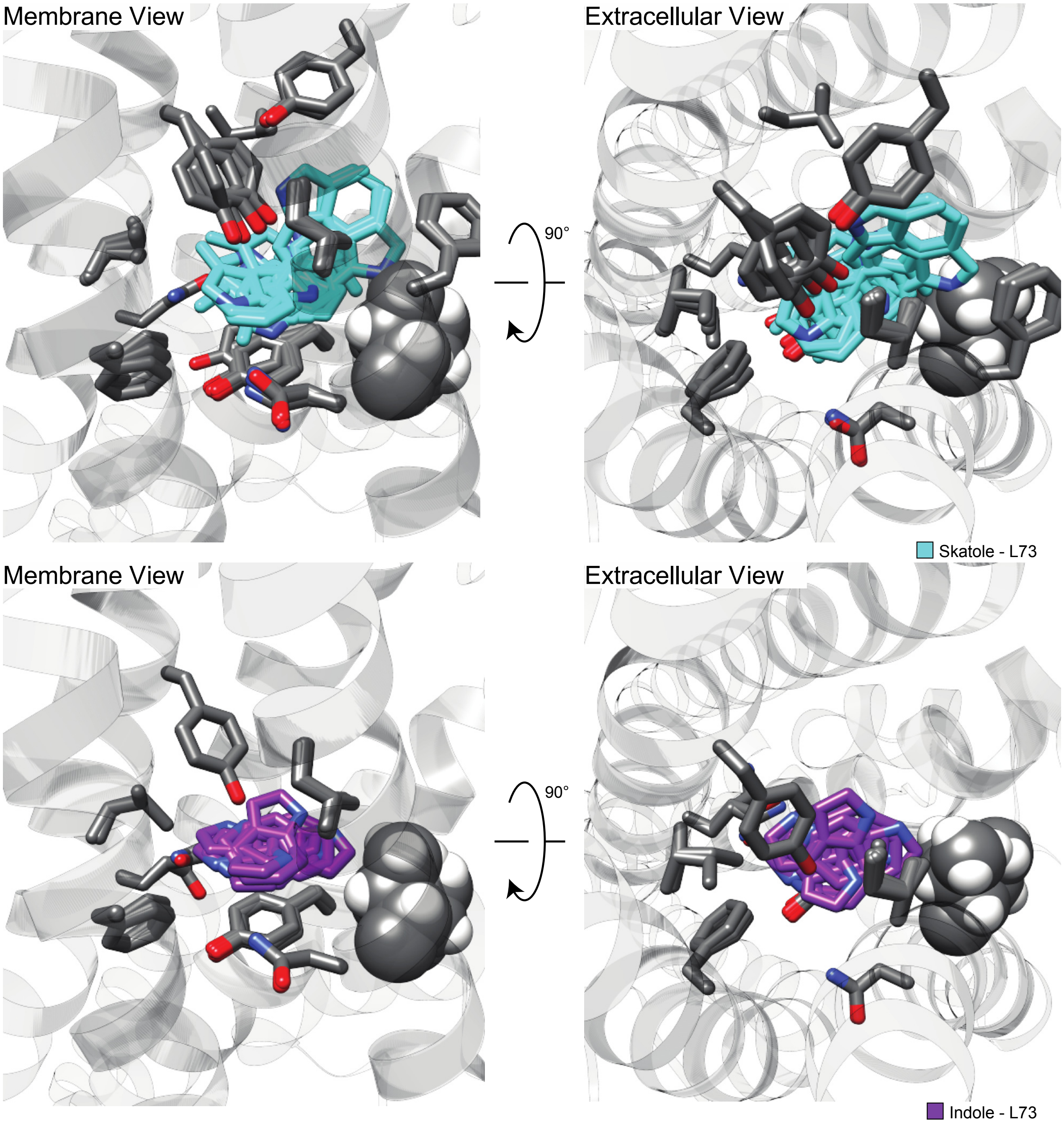


**Figure 6—supplement figure 4B.** Superposition of all skatole or indole clusters docked to CquiOR10-A73L.

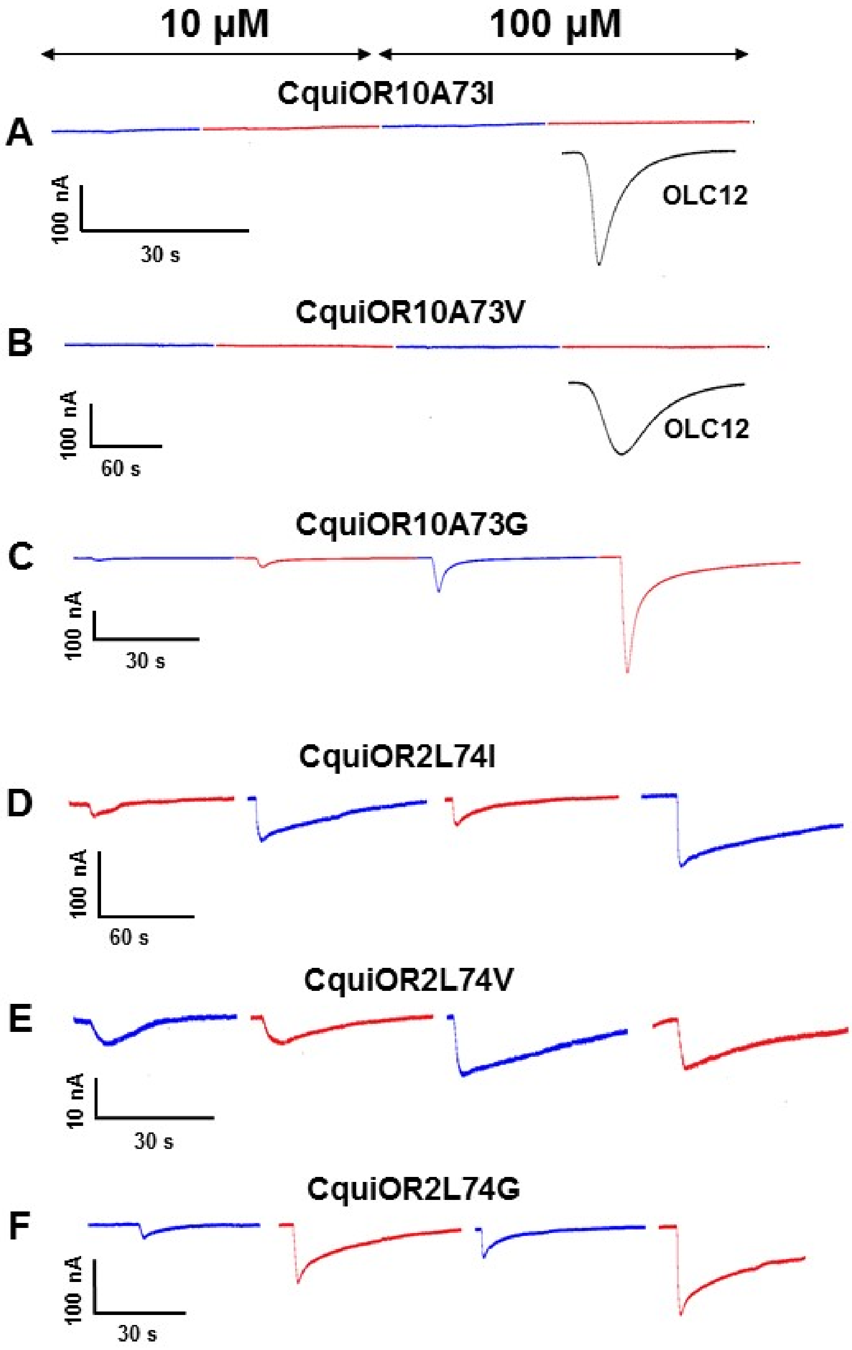


**Figure 7—supplement figure 1.** Representative traces of CquiOR10 and CquiOR2 single-point mutants. After coexpression with CquiOrco, oocytes were stimulated with skatole (red) and indole (blue) at 10 and 100 µM, and OCL12 at 100 µM. (A-C): CquiOR10 mutants; (D-F) CquiOR2 mutants.

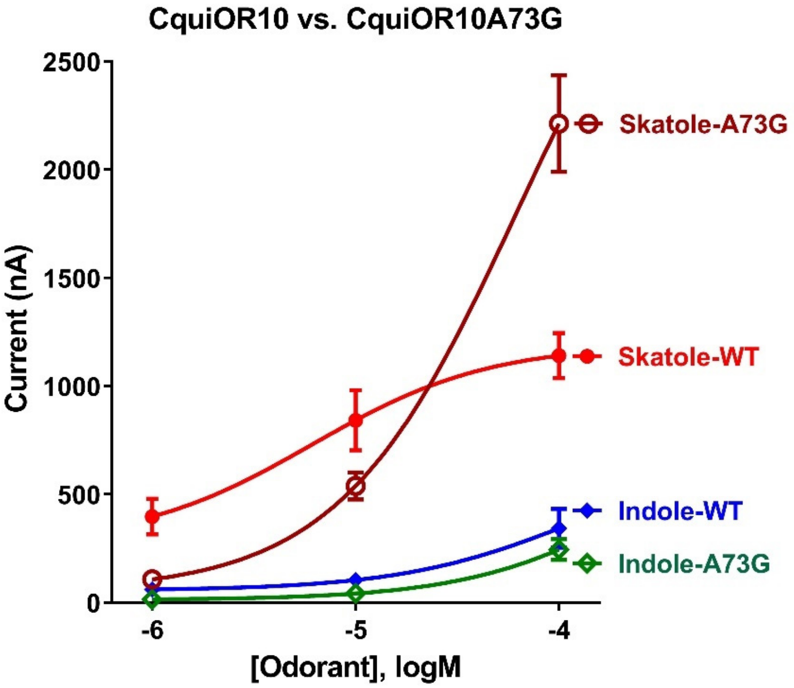


**Figure 7—supplement figure 2.** Concentration-response analysis for activation of the wildtype receptor CquiOR10 and CquiOR10A73G with skatole and indole. (For each odorant, n = 5 and 10, respectively).

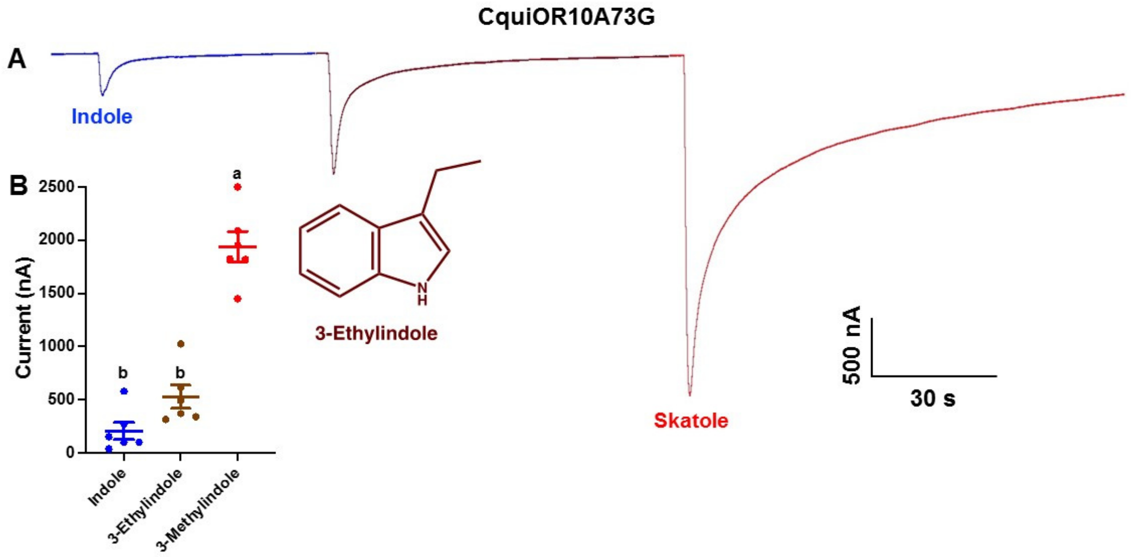


**Figure 7—supplement figure 3.** Response of CquiOR10A73G/CquiOrco-expressing oocyte to indole (blue), skatole (red), and 3-ethylindole (brown). (A) Representative trace recorder after challenging an oocyte with the three odorants at the same dose (100 μM). (B) Quantification of responses from six different oocytes. Columns with the same letter are not significantly different (Repeated measures, one-way ANOVA).

Extended—Table 1. Responses of CquiOR10Mx/CquiOrco-expressing oocytes to skatole and indole

| Chimeric OR | The Most Potent Ligand | Sensitivity Relative to CquiOR10 |
|---|---|---|
| WT | Skatole | ++ |
| M1,M2,M3,M4,M5,M6,M7 | No Response | - |
| M1,M2,M3,M4,M5,M6 | No Response | - |
| M1 | No Response | - |
| M2 | No Response | - |
| M3 | Skatole | + |
| M4 | Skatole | + |
| M5 | Skatole | +++ |
| M6 | Skatole | +++ |
| M7 | Skatole | ++ |
| M3,4,5,6,7 | Skatole | ++ |
| M3,4,5,6 | Skatole | + |
| M4,5,6 | Skatole | + |
| M3,5,6 | Skatole | + |
| M3,4,6 | Skatole | + |
| M3,4,5 | Skatole | + |
| M3,4 | Skatole | ++ |
| M3,5 | Skatole | +++ |
| M3,6 | Skatole | + |
| M3,7 | Skatole | ++ |
| M4,5 | Skatole | +++ |
| M4,6 | Skatole | ++ |
| M4,7 | Skatole | ++ |
| M5,6 | Skatole | ++ |
| M5,7 | Skatole | +++ |
| M6,7 | Skatole | +++ |
| M1,2 | No Response | - |
| M1,3 | No Response | - |
| M1,4 | No Response | - |
| M1,5 | No Response | - |
| M1,6 | No Response | - |
| M1,7 | No Response | - |
| M2,3 | No Response | - |
| M2,4 | No Response | - |
| M2,5 | No Response | - |
| M2,6 | No Response | - |
| M2,7 | Indole | + |

Extended—Table 2. Lowest Rosetta interface score and lowest RMSD of 10 largest eugenol clusters docked to MhraOR5.[†]

| | **Lowest interface score (REU)** | **RMSD (Å)** | **Cluster size** |
|---|---|---|---|
| Cluster 1 | -17.8 | 0.75 | 1875 |
| Cluster 2 | -18.1 | 4.96 | 1283 |
| Cluster 3 | -19.2 | 5.05 | 912 |
| Cluster 4 | -17.6 | 2.79 | 860 |
| Cluster 5 | -18.3 | 2.82 | 665 |
| Cluster 6 | -17.9 | 2.93 | 656 |
| Cluster 7 | -17.8 | 2.79 | 563 |
| Cluster 8 | -18.3 | 0.55 | 510 |
| Cluster 9 | -21.2 | 2.37 | 322 |
| Cluster 10 | -17.9 | 2.82 | 305 |

[†] Values should not be compared across columns since they originate from different predictions; values should only be compared within a column. REU: Rosetta Energy Units.

Extended—Table 3. Lowest Rosetta interface score of 10 largest skatole clusters docked to CquiOR10 and CquiOR10-A73L

| | **CquiOR10** | | **CquiOR10-A73L** | |
|---|---|---|---|---|
| | Lowest interface score (REU) | Cluster size | Lowest interface score (REU) | Cluster size |
| Cluster 1 | -12.6 | 661 | -14.6 | 539 |
| Cluster 2 | -12.0 | 419 | -13.5 | 502 |
| Cluster 3 | -13.8 | 360 | -14.1 | 432 |
| Cluster 4 | -13.1 | 350 | -13.8 | 424 |
| Cluster 5 | -11.4 | 301 | -15.6 | 405 |
| Cluster 6 | -12.9 | 291 | -13.6 | 385 |
| Cluster 7 | -12.9 | 290 | -13.3 | 369 |
| Cluster 8 | -13.0 | 284 | -12.3 | 360 |
| Cluster 9 | -13.4 | 269 | -13.3 | 336 |
| Cluster 10 | -12.8 | 262 | -13.2 | 321 |

Extended—Table 4. Lowest Rosetta interface score of 10 largest indole clusters docked to CquiOR10 and CquiOR10-A73L.

| | **CquiOR10** | | **CquiOR10-A73L** | |
|---|---|---|---|---|
| | Lowest interface score (REU) | Cluster size | Lowest interface score (REU) | Cluster size |
| Cluster 1 | -12.6 | 607 | -13.0 | 596 |
| Cluster 2 | -13.0 | 472 | -12.9 | 575 |
| Cluster 3 | -11.9 | 376 | -12.4 | 494 |
| Cluster 4 | -11.4 | 341 | -11.8 | 467 |
| Cluster 5 | -11.9 | 341 | -13.7 | 400 |
| Cluster 6 | -11.0 | 318 | -12.4 | 387 |
| Cluster 7 | -11.2 | 299 | -11.4 | 340 |
| Cluster 8 | -10.6 | 282 | -11.6 | 309 |
| Cluster 9 | -11.0 | 264 | -11.5 | 304 |
| Cluster 10 | -10.4 | 237 | -11.5 | 289 |

Extended—Table 5. Hydrophobic interactions reported by the lowest interface-energy prediction from each cluster. The total PLIP-reported interactions† and the total unique residues forming hydrophobic interactions with skatole/indole are reported.

| Cluster | CquiOR10 — skatole all PLIP-reported contacts | CquiOR10 — skatole unique residues | CquiOr10-A73L — skatole all PLIP-reported contacts | CquiOr10-A73L — skatole unique residues | CquiOR10 — indole all PLIP-reported contacts | CquiOR10 — indole unique residues | CquiOr10-A73L — indole all PLIP-reported contacts | CquiOr10-A73L — indole unique residues |
|---|---|---|---|---|---|---|---|---|
| 1 | 8 | 5 | 5 | 5 | 4 | 4 | 4 | 4 |
| 2 | 6 | 5 | 6 | 6 | 4 | 4 | 5 | 4 |
| 3 | 8 | 7 | 4 | 4 | 5 | 5 | 5 | 5 |
| 4 | 6 | 5 | 6 | 6 | 5 | 5 | 3 | 3 |
| 5 | 6 | 4 | 5 | 5 | 3 | 2 | 5 | 4 |
| 6 | 5 | 3 | 5 | 5 | 6 | 5 | 5 | 5 |
| 7 | 3 | 3 | 7 | 5 | 6 | 5 | 2 | 2 |
| 8 | 8 | 6 | 4 | 4 | 3 | 2 | 3 | 3 |
| 9 | 7 | 6 | 5 | 5 | 6 | 3 | 5 | 5 |
| 10 | 6 | 5 | 5 | 4 | 4 | 3 | 2 | 2 |
| Median | 6.0 | 5.0 | 5.0 | 5.0 | 4.5 | 4.0 | 4.5 | 4.0 |
| Mean | 6.3 | 4.9 | 5.2 | 4.9 | 4.6 | 3.8 | 3.9 | 3.7 |
| Standard Deviation | 1.6 | 1.3 | 0.9 | 0.7 | 1.2 | 1.2 | 1.3 | 1.2 |

† PLIP does not report the total number of hydrophobic interactions. Rather, the total hydrophobic interactions are clustered to 1) report interactions that are not π-stacking, 2) report the closest interaction distance a ligand atom interacts with binding site atoms, and 3) report the closest interaction distance a protein atom interacts with neighboring ligand atoms.

Extended—Table 6. Lowest interface-energy predictions with Leu-73 hydrophobic interactions reported by PLIP.†

| Cluster | CquiOr10-A73L — skatole | CquiOr10-A73L — indole |
|---|---|---|
| **2** | 1 | -- |
| **4** | 1 | -- |
| **5** | 1 | 1 |
| **6** | -- | 1 |
| **7** | 1 | -- |
| **9** | 1 | -- |

† The lowest interface-energy prediction was used to represent each cluster.

Extended—Table 7. Total PLIP-reported[†] hydrophobic interactions* reported per-residue with indole/skatole.

| Residue | CquiOR10 — skatole | CquiOr10-A73L — skatole | CquiOR10 — indole | CquiOr10-A73L — indole | Mean skatole predictions | Mean indole predictions | Mean all structure predictions |
|---|---|---|---|---|---|---|---|
| **Phe-66** | -- | -- | 1 | -- | -- | 1 | 1 |
| **Ile-69** | 7 | 9 | 7 | 9 | 8 | 8 | 8 |
| **Phe-70** | 1 | 2 | 1 | -- | 2 | 1 | 1 |
| **Asn-72** | 7 | 5 | 5 | 4 | 6 | 4 | 5 |
| **Ala/Leu73** | -- | 5 | -- | 2 | 5 | 2 | 4 |
| **Ile-134** | 1 | 1 | -- | -- | 1 | -- | 1 |
| **Tyr-138** | -- | 2 | 2 | -- | 2 | 2 | 2 |
| **Tyr-152** | 1 | -- | -- | -- | 1 | -- | 1 |
| **Tyr-185** | 10 | 10 | 8 | 10 | 10 | 9 | 10 |
| **Ile-186** | -- | -- | 2 | -- | -- | 2 | 2 |
| **Phe-261** | 6 | 5 | 6 | 5 | 6 | 6 | 6 |
| **Tyr-287** | 8 | 6 | 2 | 3 | 7 | 2 | 5 |
| **Ile-291** | 4 | 4 | 1 | 2 | 4 | 2 | 3 |
| **Gln-294** | 3 | -- | 3 | 2 | 3 | 2 | 3 |

[†] PLIP does not report the total number of hydrophobic interactions. Rather, the total hydrophobic interactions are clustered to 1) report interactions that are not π-stacking, 2) report the closest interaction distance a ligand atom interacts with binding site atoms, and 3) report the closest interaction distance a protein atom interacts with neighboring ligand atoms.
* Values represent the total number of representative predictions interacting with the specified residue. Each cluster is represented by the lowest interface-energy prediction.

Extended—Table 8. Hydrogen bond interactions reported by the lowest interface-energy prediction from each cluster. The total PLIP-reported interactions† and the total unique residues forming hydrogen bonds with skatole/indole are reported.

| Cluster | CquiOR10 — skatole all PLIP-reported contacts | CquiOR10 — skatole unique residues | CquiOr10-A73L — skatole all PLIP-reported contacts | CquiOr10-A73L — skatole unique residues | CquiOR10 — indole all PLIP-reported contacts | CquiOR10 — indole unique residues | CquiOr10-A73L — indole all PLIP-reported contacts | CquiOr10-A73L — indole unique residues |
|---|---|---|---|---|---|---|---|---|
| **1** | -- | -- | -- | -- | -- | -- | 1 | 1 |
| **2** | -- | -- | -- | -- | -- | -- | -- | -- |
| **3** | -- | -- | -- | -- | 1 | 1 | -- | -- |
| **4** | -- | -- | 1 | 1 | -- | -- | -- | -- |
| **5** | -- | -- | 1 | 1 | -- | -- | 1 | 1 |
| **6** | -- | -- | -- | -- | -- | -- | -- | -- |
| **7** | 2 | 1 | -- | -- | 2 | 1 | -- | -- |
| **8** | 2 | 1 | -- | -- | 1 | 1 | -- | -- |
| **9** | -- | -- | -- | -- | 1 | 1 | -- | -- |
| **10** | -- | -- | -- | -- | -- | -- | -- | -- |
| **Median** | 0.0 | 0.0 | 0.0 | 0.0 | 0.0 | 0.0 | 0.0 | 0.0 |
| **Mean** | 0.4 | 0.2 | 0.2 | 0.2 | 0.5 | 0.4 | 0.2 | 0.2 |
| **Standard Deviation** | 0.8 | 0.4 | 0.4 | 0.4 | 0.7 | 0.5 | 0.4 | 0.4 |

† Hydrogen bond interactions reported by PLIP were filtered using previously reported bond distances (Bissantz et al., 2010).

Extended—Table 9. Total PLIP-reported† hydrogen bond interactions reported per-residue with indole/skatole.

| Residue | CquiOR10 — skatole | CquiOr10-A73L — skatole | CquiOR10 — indole | CquiOr10-A73L — indole | Mean skatole predictions | Mean indole predictions | Mean all structure predictions |
|---|---|---|---|---|---|---|---|
| **Ile-69** | -- | -- | 1 | -- | -- | 1 | 1 |
| **Asn-72** | -- | 1 | -- | -- | 1 | -- | 1 |
| **Tyr-138** | -- | -- | 1 | -- | -- | 1 | 1 |
| **Tyr-287** | -- | -- | 1 | -- | -- | 1 | 1 |
| **Gln-294** | 2 | 1 | 1 | 2 | 2 | 2 | 2 |

† Hydrogen bond interactions reported by PLIP were filtered using previously reported bond distances(Bissantz et al., 2010).

Extended—Table 10. Descriptions of each hydrogen bond reported by each cluster's representative† prediction.

| Structure — Ligand | Cluster | Residue | Donor* | Acceptor* | Donor-Acceptor Distance (Å) |
|---|---|---|---|---|---|
| **CquiOR10 — skatole** | 7 | Gln-294 | Gln-294 sidechain, Nam | Skatole, Npl | 3.1 |
| **CquiOR10 — skatole** | 7 | Gln-294 | Skatole, Npl | Gln-294 sidechain, O2 | 2.9 |
| **CquiOR10 — skatole** | 8 | Gln-294 | Gln-294 sidechain, Nam | Skatole, Npl | 3.1 |
| **CquiOR10 — skatole** | 8 | Gln-294 | Skatole, Npl | Gln-294 sidechain, O2 | 2.7 |
| **CquiOR10-A73L — skatole** | 4 | Asn-72 | Asn-72 sidechain, Nam | Skatole, Npl | 3.2 |
| **CquiOR10-A73L — skatole** | 5 | Gln-294 | Skatole, Npl | Gln-294 sidechain, O2 | 2.8 |
| **CquiOR10 — indole** | 3 | Tyr-287 | Tyr-287 sidechain, O3 | Indole, Npl | 3.0 |
| **CquiOR10 — indole** | 7 | Gln-294 | Gln-294 sidechain, Nam | Indole, Npl | 3.1 |
| **CquiOR10 — indole** | 7 | Gln-294 | Indole, Npl | Gln-294 sidechain, O2 | 2.8 |
| **CquiOR10 — indole** | 8 | Ile-69 | Indole, Npl | Ile-69 backbone, O2 | 3.1 |
| **CquiOR10 — indole** | 9 | Tyr-138 | Indole, Npl | Tyr-138 sidechain, O3 | 2.8 |
| **CquiOR10-A73L — indole** | 1 | Gln-294 | Indole, Npl | Gln-294 sidechain, O2 | 2.9 |
| **CquiOR10-A73L — indole** | 5 | Gln-294 | Indole, Npl | Gln-294 sidechain, O2 | 3.0 |

† The lowest interface-energy prediction was used to represent each cluster.
* The atom type definition is derived from IDATM nomenclature (Meng & Lewis, 1991).

Extended—Table 11. Parallel pi-stacking interactions reported by the lowest interface-energy prediction from each cluster. The total PLIP-reported interactions† and the total unique residues forming parallel pi-stacking interactions with skatole/indole are reported.

| Cluster | CquiOR10 — skatole all PLIP-reported contacts | CquiOR10 — skatole unique residues | CquiOr10-A73L — skatole all PLIP-reported contacts | CquiOr10-A73L — skatole unique residues | CquiOR10 — indole all PLIP-reported contacts | CquiOR10 — indole unique residues | CquiOr10-A73L — indole all PLIP-reported contacts | CquiOr10-A73L — indole unique residues |
|---|---|---|---|---|---|---|---|---|
| **1** | -- | -- | 1 | 1 | -- | -- | 1 | 1 |
| **2** | -- | -- | -- | -- | -- | -- | 1 | 1 |
| **3** | -- | -- | -- | -- | -- | -- | 1 | 1 |
| **4** | -- | -- | 1 | 1 | -- | -- | -- | -- |
| **5** | -- | -- | -- | -- | 2 | 2 | 1 | 1 |
| **6** | -- | -- | -- | -- | -- | -- | -- | -- |
| **7** | -- | -- | -- | -- | -- | -- | -- | -- |
| **8** | -- | -- | -- | -- | -- | -- | 1 | 1 |
| **9** | -- | -- | -- | -- | -- | -- | 1 | 1 |
| **10** | -- | -- | -- | -- | -- | -- | 2 | 1 |
| **Median** | 0.0 | 0.0 | 0.0 | 0.0 | 0.0 | 0.0 | 1.0 | 1.0 |
| **Mean** | 0.0 | 0.0 | 0.2 | 0.2 | 0.2 | 0.2 | 0.8 | 0.7 |
| **Standard Deviation** | 0.0 | 0.0 | 0.4 | 0.4 | 0.6 | 0.6 | 0.6 | 0.5 |

† Parallel pi-stacking interactions were filtered by previously reported bond distances (Bissantz et al., 2010). No perpendicular pi-stacking interactions met filtering distance criteria.

Extended—Table 12. Total PLIP-reported parallel pi-stacking interactions† reported per-residue with indole/skatole.

| Residue | CquiOR10 — skatole | CquiOr10-A73L — skatole | CquiOR10 — indole | CquiOr10-A73L — indole | Mean skatole predictions | Mean indole predictions | Mean all structure predictions |
|---|---|---|---|---|---|---|---|
| **Tyr-138** | -- | -- | 1 | -- | -- | 1 | 1 |
| **Tyr-152** | -- | -- | 1 | -- | -- | 1 | 1 |
| **Tyr-185** | -- | 2 | -- | 8 | 2 | 8 | 5 |

† Parallel pi-stacking interactions were filtered by previously reported bond distances(Bissantz et al., 2010). No perpendicular pi-stacking interactions met filtering distance criteria.

Extended—Table 13. Descriptions of each parallel pi-stacking interaction reported by each cluster's representative† prediction.

| Structure — Ligand | Cluster | Residue | Central ring distance (Å)* |
|---|---|---|---|
| **CquiOR10-A73L — skatole** | 1 | Tyr-185 | 3.6 |
| **CquiOR10-A73L — skatole** | 4 | Tyr-185 | 3.4 |
| **CquiOR10 — indole** | 5 | Tyr-138 | 3.5 |
| **CquiOR10 — indole** | 5 | Tyr-152 | 3.4 |
| **CquiOR10-A73L — indole** | 1 | Tyr-185 | 3.4 |
| **CquiOR10-A73L — indole** | 2 | Tyr-185 | 3.6 |
| **CquiOR10-A73L — indole** | 3 | Tyr-185 | 3.6 |
| **CquiOR10-A73L — indole** | 5 | Tyr-185 | 3.4 |
| **CquiOR10-A73L — indole** | 8 | Tyr-185 | 3.6 |
| **CquiOR10-A73L — indole** | 9 | Tyr-185 | 3.6 |
| **CquiOR10-A73L — indole** | 10 | Tyr-185 | 3.6 |
| **CquiOR10-A73L — indole** | 10 | Tyr-185 | 3.6 |

† The lowest interface-energy prediction was used to represent each cluster.
* The central distance is computed by two centroids positioned at the center each aromatic ring.

Extended—Table 14. Cartesian coordinates of C-alpha atoms at position 73 from CquiOR10/A73L docking. The lowest interface-scoring model was used from each cluster. Models were aligned to helix S7b prior to recording position.

| | Skatole | | | | | | Indole | | | | | |
|---|---|---|---|---|---|---|---|---|---|---|---|---|
| | CquiOR10 | | | CquiOr10-A73L | | | CquiOR10 | | | CquiOr10-A73L | | |
| | x | y | z | x | y | z | x | y | z | x | y | z |
| Cluster 1 | 164.917 | 184.462 | 155.034 | 165.788 | 183.953 | 155.051 | 164.914 | 184.490 | 155.036 | 165.797 | 183.967 | 155.054 |
| Cluster 2 | 164.936 | 184.513 | 155.026 | 165.801 | 183.981 | 155.051 | 164.928 | 184.470 | 155.066 | 165.808 | 183.975 | 155.054 |
| Cluster 3 | 164.934 | 184.465 | 155.063 | 165.795 | 183.961 | 155.057 | 164.967 | 184.470 | 155.012 | 165.800 | 183.964 | 155.058 |
| Cluster 4 | 164.925 | 184.485 | 155.038 | 165.807 | 183.970 | 155.061 | 164.920 | 184.464 | 155.052 | 165.796 | 183.976 | 155.055 |
| Cluster 5 | 164.934 | 184.436 | 155.038 | 165.885 | 184.086 | 155.111 | 164.938 | 184.488 | 155.016 | 165.872 | 184.010 | 155.119 |
| Cluster 6 | 164.914 | 184.533 | 155.032 | 165.790 | 184.023 | 155.039 | 164.917 | 184.541 | 155.034 | 165.803 | 183.971 | 155.048 |
| Cluster 7 | 164.953 | 184.446 | 155.039 | 165.795 | 183.972 | 155.046 | 164.940 | 184.472 | 155.049 | 165.791 | 183.968 | 155.060 |
| Cluster 8 | 164.927 | 184.477 | 155.043 | 165.776 | 183.949 | 155.063 | 164.937 | 184.489 | 155.034 | 165.870 | 184.015 | 155.113 |
| Cluster 9 | 164.923 | 184.487 | 155.036 | 165.915 | 184.044 | 155.142 | 164.935 | 184.534 | 155.035 | 165.792 | 183.957 | 155.051 |
| Cluster 10 | 164.931 | 184.439 | 155.050 | 165.797 | 184.009 | 155.058 | 164.932 | 184.442 | 155.045 | 165.790 | 183.956 | 155.052 |
| Mean | 164.929 | 184.474 | 155.040 | 165.815 | 183.995 | 155.068 | 164.933 | 184.486 | 155.038 | 165.812 | 183.976 | 155.066 |
| Standard Deviation (Å) | 0.011 | 0.032 | 0.010 | 0.0461 | 0.045 | 0.032 | 0.015 | 0.031 | 0.016 | 0.032 | 0.020 | 0.026 |

Extended—Table 15. Mean distances (Å) of C-alpha atoms at position 73 from CquiOR10/A73L docking using skatole and indole. The lowest interface-scoring model was used from each cluster. Models were aligned to helix S7b prior to distance calculation.

| **Protein — Ligand** | **CquiOR10 — skatole** | **CquiOr10-A73L — skatole** | **CquiOR10 — indole** | **CquiOr10-A73L — indole** |
|---|---|---|---|---|
| **CquiOR10 — skatole** | -- | -- | -- | -- |
| **CquiOr10-A73L — skatole** | 1.007 | -- | -- | -- |
| **CquiOR10 — indole** | 0.012 | 1.010 | -- | -- |
| **CquiOr10-A73L — indole** | 1.014 | 0.019 | 1.017 | -- |
